# The phenological response of European vegetation to urbanisation is mediated by macrobioclimatic factors

**DOI:** 10.1101/2023.05.31.543030

**Authors:** Javier Galán Díaz, Adela Montserrat Gutiérrez-Bustillo, Jesús Rojo

## Abstract

Plant phenology is a crucial component of ecosystem functioning and is affected by multiple elements of global change; we therefore need to quantify the current phenological changes associated to human activities and understand their impacts on ecosystems. Urbanisation and the intensification of anthropogenic activities alter meteorological conditions and cause phenological changes in urban vegetation worldwide. We used remote sensing data to evaluate the phenological response (start of season date SOS, length of season LOS and end of season date EOS) to urbanisation in the 69 most populated pan-European metropolitan areas (i.e., those that include cities with a population over 450,000 inhabitants) for the period 2002-2021, taking into account the main vegetation types (evergreen forests, deciduous forests, mixed forests, sparse woody vegetation and grasslands). We found that macrobioclimatic factors strongly determined the strength and direction of the phenological response to urbanisation intensity. In general, SOS advanced and LOS increased with urbanisation intensity across Europe. However, the greatest advances in SOS were registered in metropolitan areas in the Mediterranean region, where there was also more uncertainty in the relationship between SOS and urbanisation. The EOS advanced with urbanisation in metropolitan areas in the Mediterranean macrobioclimate (except for deciduous forests), whereas the opposite trend was observed in metropolitan areas from the Temperate macrobioclimate. Our results suggest that macrobioclimatic constraints operating at the continental scale mediate the relationship between plant phenology and urbanisation intensity.

## Introduction

Plant phenology is a crucial component of ecosystem functioning because of its role in the carbon cycle (Piao et al., 2023), feedbacks on local and regional climate (Richardson et al., 2013) and the assembly of ecological communities (Inouye, 2022; Renner and Zohner, 2018). It also has important repercussions from the point of view of public-health as is the case of the flowering phenology of species with high allergenic potential (Bernard-Verdier et al., 2022; Rojo et al., 2021). Plant phenology is sensitive to nutrient and water availability and changes in temperature and photoperiod (Piao et al., 2023); accordingly, over the last decades scientists have been making great efforts to quantify phenological changes in response to the multiple components of global change and to understand their consequences on ecosystems (Garcia et al., 2014; Menzel et al., 2020).

The intensification of urbanisation and anthropogenic activities alter meteorological conditions at the local scale and affect the dynamics of urban vegetation (Trusilova and Churkina, 2008). Higher air and land surface temperatures in cities compared to the surrounding rural areas – a phenomenon referred to as the urban heat island – is the most prominent driver of phenological changes associated to urbanisation (Jochner and Menzel, 2015; Meng et al., 2020; Neil and Wu, 2006). Plants’ phenological response is also affected by other meteorological changes derived from urbanisation such as a decrease in air humidity (Hao et al., 2018), and stressors such as limited space for growth, soil compaction and pollution (Czaja et al., 2020).

Data gathered from ground stations and satellite observations have made it possible to unveil the general responses of plant phenology to urbanisation. Most studies agree that spring phenophases (i.e., flowering, leaf-out and bud opening) tend to start earlier in highly urbanised areas (Jia et al., 2021; Li et al., 2019; Meng et al., 2020; Ren et al., 2018; Wohlfahrt et al., 2019; Yang et al., 2020). The opposite trend – i.e., delay in spring phenophases with urbanisation – has occasionally been reported (Galán Díaz et al., 2023; Zhang et al., 2022), which may indicate that some plant species do not adequately meet their chilling requirements due to rising autumn and winter temperatures (Inouye, 2022; Piao et al., 2023; Yu et al., 2010). Most studies also report delays in autumn phenophases (e.g., leaf senescence) caused by urbanisation (Ren et al., 2018; Wohlfahrt et al., 2019; Yang et al., 2020). However, less consistent trends have been found between urbanisation intensity and autumn phenophases, probably because they also are highly dependent on other factors such as photoperiod, preseason forcing and productivity rates during the growth season (Keenan and Richardson, 2015; Wu et al., 2018; Zani et al., 2020).

The direction and strength of the relationship between plant phenology and urbanisation is affected by other factors such as plant functional type and regional climate. On the one hand, the response of different vegetation types to urbanisation is mediated by temperature sensitivity and physiological constraints. For instance, annual plants, early-flowering species and lower-growing perennials with a higher specific leaf area show the greatest advances in their spring phenophases (Inouye, 2022; Li et al., 2019; Neil and Wu, 2006). In addition, it has been reported that deciduous and evergreen species might show contrasting phenological responses to urbanisation; hence leaf longevity also plays an important role (Galán Díaz et al., 2023; Zhang et al., 2022). On the other hand, regional climate affects the daily and seasonal relationship between urbanisation intensity and land surface temperatures (Sismanidis et al., 2022; Wienert and Kuttler, 2005). For instance, in mainland China, Ma et al. (2016) reported a stronger relationship between urbanisation and summer land surface temperature in dry and cold regions compared to wet and hot regions. Other regional climatic constraints, such as moisture availability, mediate the plant phenological response to urbanisation (Kabano et al., 2021). We might therefore expect plants’ phenological response to urbanisation to depend on regional climate.

Recent efforts to understand the effect of urbanisation on plant phenology at the species-scale in Europe using ground observations have reported that flowering, fruiting and leaf-out dates occurred an average of 1.0-4.3 days earlier in completely urbanised compared to non-urbanised areas (Li et al., 2019; Wohlfahrt et al., 2019); and that leaf senescence tended to occur 1.3–2.7 days later (Wohlfahrt et al., 2019). These studies provided invaluable information on the response of several phenophases of individual species to urbanisation, but they focused on Europe’s Temperate region (Templ et al., 2018). We argue that the use of other data sources and methodologies will allow to further our understanding of the relationship between urbanisation and vegetation phenology in Europe. In particular, using remote sensing data, we can quantify the plant phenological response to urbanisation at the landscape-scale (i.e., main vegetation types) within metropolitan areas, and explore changes in the strength and direction of this relationship across the entire Temperate-Mediterranean gradient in Europe.

In this study we used remotely sensed data to quantify the impact of urbanisation intensity on the phenology of evergreen, deciduous and mixed forests, sparse woody vegetation (i.e., forest canopy cover of less than 60%) and grasslands in the main 69 pan-European metropolitan areas (i.e., those that include cities with a population over 450,000 inhabitants) for the period 2002-2021. We hypothesised that: (i) urbanisation brings forward the start of the season (SOS) of European vegetation, increases its length (LOS) and delays its end (EOS); (ii) vegetation types respond differently to urbanisation; and (iii) the phenological response of vegetation to urbanisation depends on regional climate and varies between the two main macrobioclimates present in Europe (Temperate vs. Mediterranean).

## Material and methods

### Study area

The study area comprises the main 69 pan-European metropolitan areas (Figure 1). Taking advantage of the fact that urbanisation data was also available for Turkey, we considered cities from the Mediterranean areas of this country in order to have a greater representation of cases from the Mediterranean macrobioclimate. The study area comprises two macrobioclimatic regions, i.e., Temperate and Mediterranean. In contrast to the Temperate, the Mediterranean macrobioclimate is characterised by having at least two consecutive dry months during the hottest period of the year (Rivas-Martínez et al., 2002). For each European city with a population over 450,000, we set a surrounding buffer that was two times the area of its urban extent using Zhao et al.’s (2022) global dataset of annual urban extents from night-time lights. This ensured an equal representation of urban, peri-urban and rural areas. Cities located within each other’s buffer area were treated as the same metropolitan area. For instance, Amsterdam, Rotterdam, The Hague and surrounding cities were merged within the metropolitan area of Randstad; and Köln, Essen, Dortmund, Düsseldorf, Duisburg and nearby cities were merged within the Rhine-Ruhr metropolitan area.

**Figure 1.**
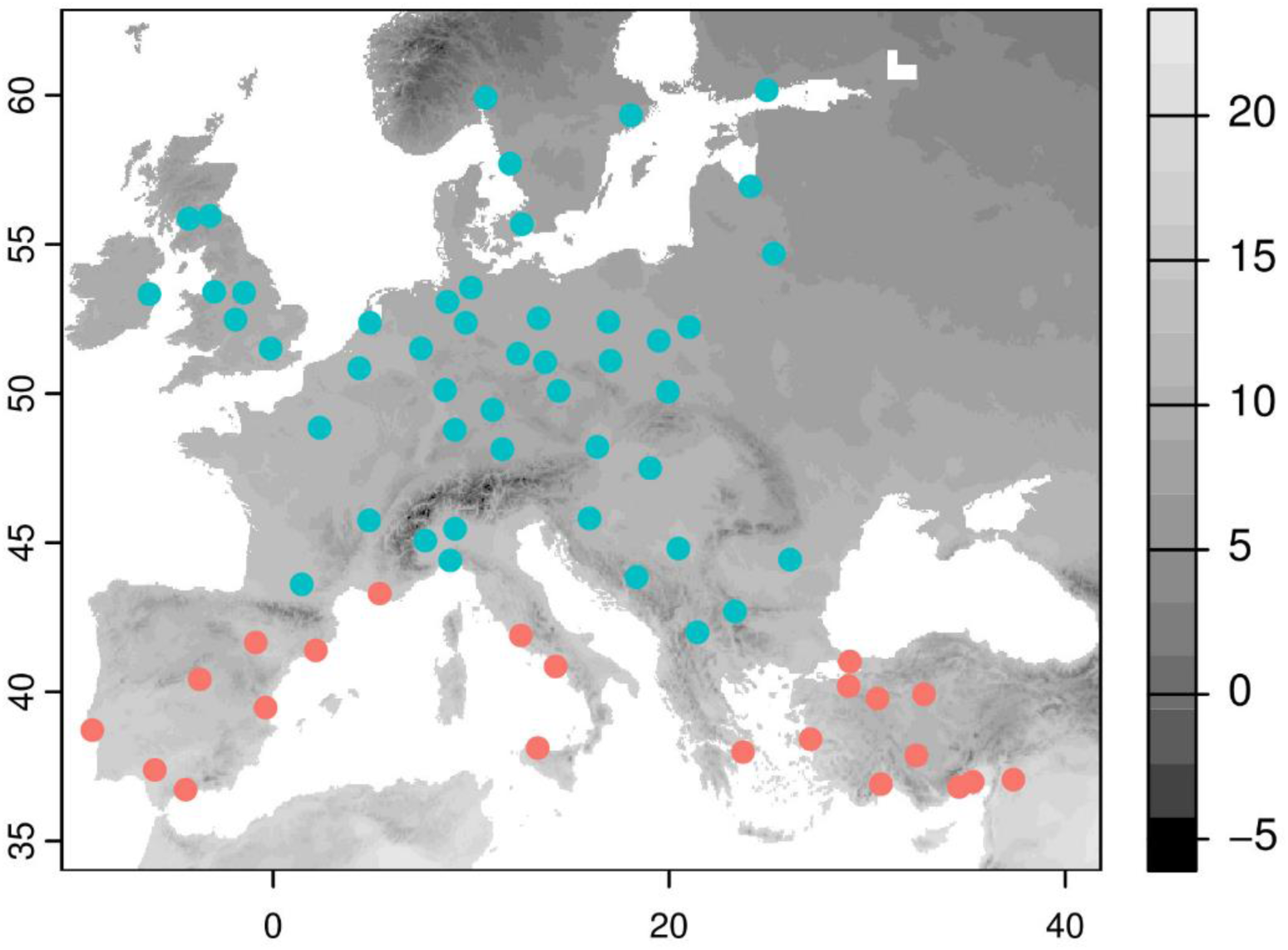
Metropolitan areas included in this study and mean annual temperature for the period 2002-2021. Metropolitan areas were classified according to their macrobioclimate in Mediterranean (red) and Temperate (blue) according to Rivas-Martínez et al. (2002).

### Data collection

Phenology and land-use data for the period 2002-2021 were obtained from MODIS and downloaded using the function *mt_subset* implemented in the MODISTools R package (Hufkens, 2022). Start of season (SOS) and end of season (EOS) dates correspond to the Greenup and Dormancy layers of the Land Cover Dynamics Yearly L3 Global 500 m dataset (MCD12Q2 v006) derived from the Enhanced Vegetation Index 2. SOS and EOS respectively are the day of the year (doy) when the vegetation index first and last crossed 15% of the segment of its amplitude. Length of season (LOS) is the number of days between SOS and EOS. Land cover data was obtained from the LC_Type1 layer of the Land Cover Type Yearly L3 Global 500 m dataset (MCD12Q1 v006) which corresponds to the Annual International Geosphere-Biosphere Programme (IGBP) classification system. We aggregated the land uses of the IGBP classification into five main vegetation types: evergreen forests (classes 1 and 2), deciduous forests (classes 3 and 4), mixed forests (class 5), sparse woody vegetation (classes 8 and 9, i.e., forest canopy cover less than 60%) and grasslands (class 10).

We used two datasets from the Copernicus Land Monitoring Service at a 100 m resolution to estimate urbanisation intensity: (i) CORINE Land Cover and (ii) Imperviousness Density. The CORINE Land Cover Inventory was available for the years 2000, 2006, 2012 and 2018. We considered urbanised areas to be those in the categories of continuous urban fabric, discontinuous urban fabric, industrial or commercial units, road and rail networks and associated land, port areas and airports. The Imperviousness Density dataset indicates the percentage of soil sealing and was available for the years 2006, 2009, 2012, 2015 and 2018. Both datasets were linearly interpolated to cover the entire period of 2002-2021 using the *approximate* function implemented in the terra R package (Hijmans et al., 2022). Extrapolation of urbanisation data was performed taking the nearest known value. For each phenological observation (i.e., centroid of each 500m^2^ pixel from MODIS), urbanisation intensity was computed as the proportion of urbanised area (i.e., CORINE dataset) and average soil sealing (i.e., Imperviousness Density) within a circular buffer with a 250m, 500m and 1km radius.

Land surface temperature (LST) was obtained from the MODIS Land Surface Temperature/Emissivity 8-Day L3 Global 1 km dataset (MOD11A2 v006) and downloaded using the function *mt_subset* implemented in the MODISTools R package (Hufkens, 2022). Elevation data was derived from the SRTM and accessed through WorldClim v2.1 at a resolution of 30 seconds (Fick and Hijmans, 2017). Climatic variables of temperature and precipitation were derived from the CRU TS monthly high-resolution gridded multivariate climate dataset (Harris et al., 2020) using the climate interpolation methods of BIOCLIM (Booth et al., 2014) and accessed through WorldClim at a resolution of 2.5 minutes (Fick and Hijmans, 2017). BIOCLIM variables for the study period were calculated using the function *biovars* implemented in the dismo R package (Hijmans et al., 2023).

### Data analysis

First, we determined the effect that the source of urbanisation intensity and buffer choice had on the analyses. For each metropolitan area, we ran six multiple regression models using SOS as the response variable, and each combination of urbanisation source (i.e., CORINE Land Cover and Imperviousness Density) and buffer size (250m, 500m and 1km radius) as the explanatory variable, including elevation and year as covariables. We then computed the Pearson coefficient among the regression coefficients of urbanisation (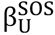) obtained from the different models. There was a high linear correlation among the phenological responses of SOS to the different proxies of urbanisation intensity (Appendix A). We used CORINE Land Cover at 500 m for subsequent analyses, which is also supported by a previous study (Wohlfahrt et al., 2019).

Second, for each metropolitan area, we used two statistical approaches to analyse the phenological response of the five main vegetation types to urbanisation. (i) We implemented a random forest model using the *randomForest* R package (Liaw & Wiener, 2002) and used partial dependent plots provided by the *pdp* R package (Greenwell, Brandon, 2017) to explore the general patterns across Europe. To show the partial effect of the urbanisation intensity on the phenological patterns, the phenological indexes were standardised by subtracting the mean and dividing by the standard deviation. This makes all phenological patterns comparable independently of the climatic conditions of each metropolitan area. (ii) We performed multiple linear regressions to quantify the effect (i.e., regression coefficients) in each city. We used the phenological indexes (SOS, LOS and EOS) as the response variables and urbanisation, year and elevation as explanatory variables (Ziello et al., 2009). We only performed multiple linear regressions in metropolitan areas where there was a minimum of 50 records per vegetation type along the time series.

Third, we explored whether the phenological response to urbanisation in European metropolitan areas depended on regional climate. This was done by performing simple linear regressions with the phenological response to urbanisation (i.e., 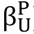) as the response variable and each of the 19 BIOCLIM variables as predictors. We then focused on the effect of annual mean temperature because it is known to influence the urban heat island (Ma et al., 2016; Wienert and Kuttler, 2005) and the phenological response of plants to urbanisation (Li et al., 2019). We then compared the phenological response of the five main vegetation types to urbanisation between the metropolitan areas of Europe’s two main macrobioclimates: Temperate vs. Mediterranean (Rivas-Martínez et al., 2002). The classification of macrobioclimates relates to the earth’s main climates and biogeographical regions and provides a simple interpretation and delimitation of the main bioregions of Europe.

Fourth, we analysed the relationship between SOS and LST to explore the importance of temperature as a driver of phenological changes along the urbanisation gradient. We used the average day and night LST during winter and spring as a reference period that includes preseason and early season temperatures (Appendix B). We used elevation and year as covariables. We excluded the other two phenological indexes for this analysis because they are also very dependent on other climatic constraints (Keenan and Richardson, 2015; Wu et al., 2018; Zani et al., 2020). Finally, we ran multiple linear regressions using urbanisation intensity, year and elevation as explanatory variables to explored whether the relationship between LST and urbanisation (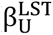) and year (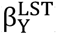) varies between metropolitan areas in the Mediterranean and Temperate macrobioclimates and depends on regional temperature.

All analyses were performed in R (version 4.3.0). The codes used in this study are available at GitHub (https://github.com/galanzse/europe_urbantrees).

## Results

The random forest model showed that, as the intensity of urbanisation rose, the start of season date (SOS) advanced and the length of season (LOS) increased in all the main vegetation types (Figure 2). Specifically, we found a significant advance in SOS with urbanisation intensity in 59.02% of cases (vegetation type × metropolitan area) when data was available, and a significant delay in 10.66% of cases (Table 1; Figure 3). Increasing urbanisation intensity resulted in a significant lengthening of the LOS in 57.38% of cases, and a shortening in 8.61%.

**Figure 2.**
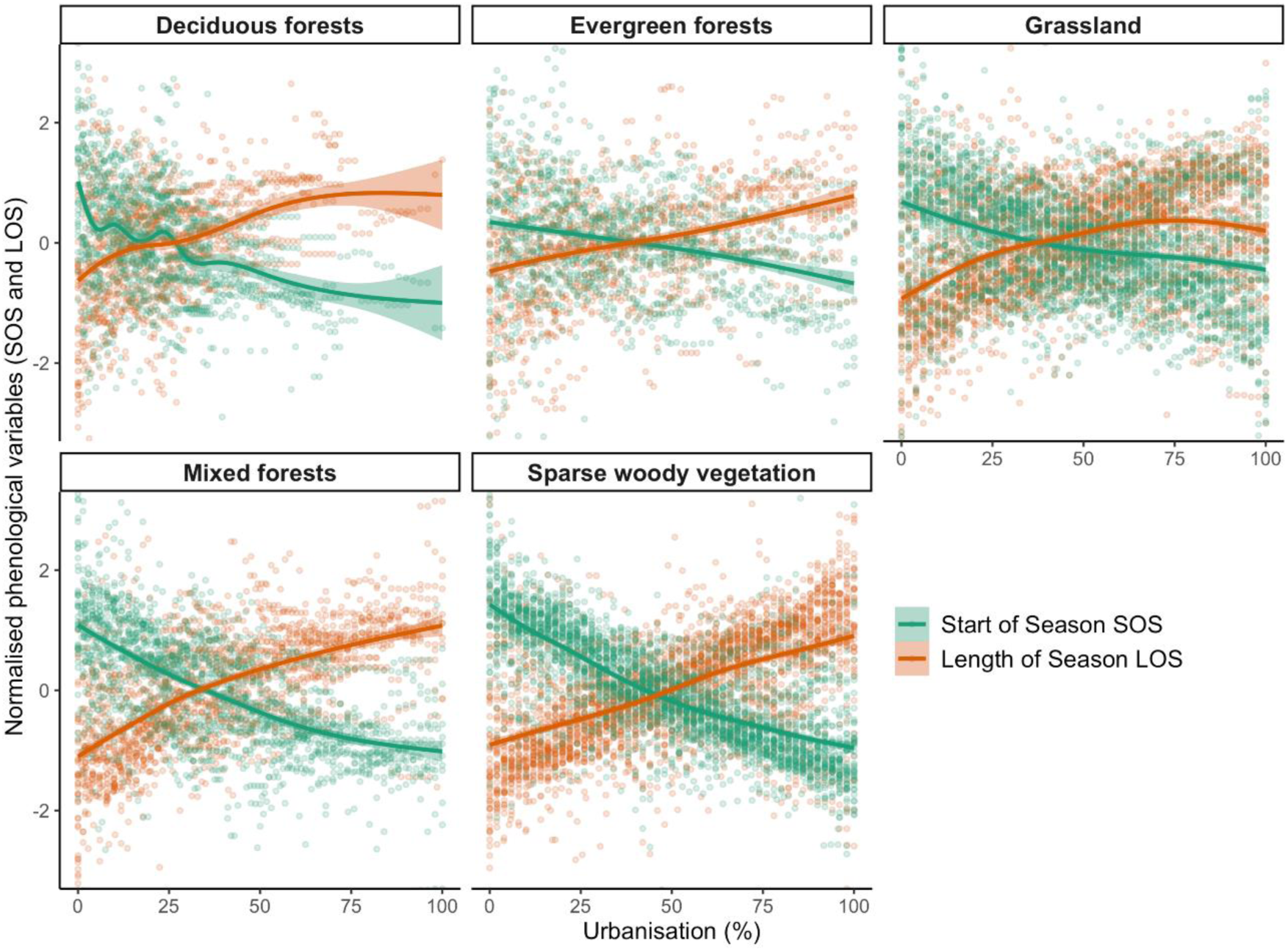
Partial effect of urbanisation intensity on the phenological indexes. Random forest models for each metropolitan area were performed to predict the general phenological response to urbanisation. The phenological variable for each metropolitan area was normalised by subtracting the mean and dividing by the standard deviation. SOS: start of season date, LOS: length of season in days.

**Figure 3.**
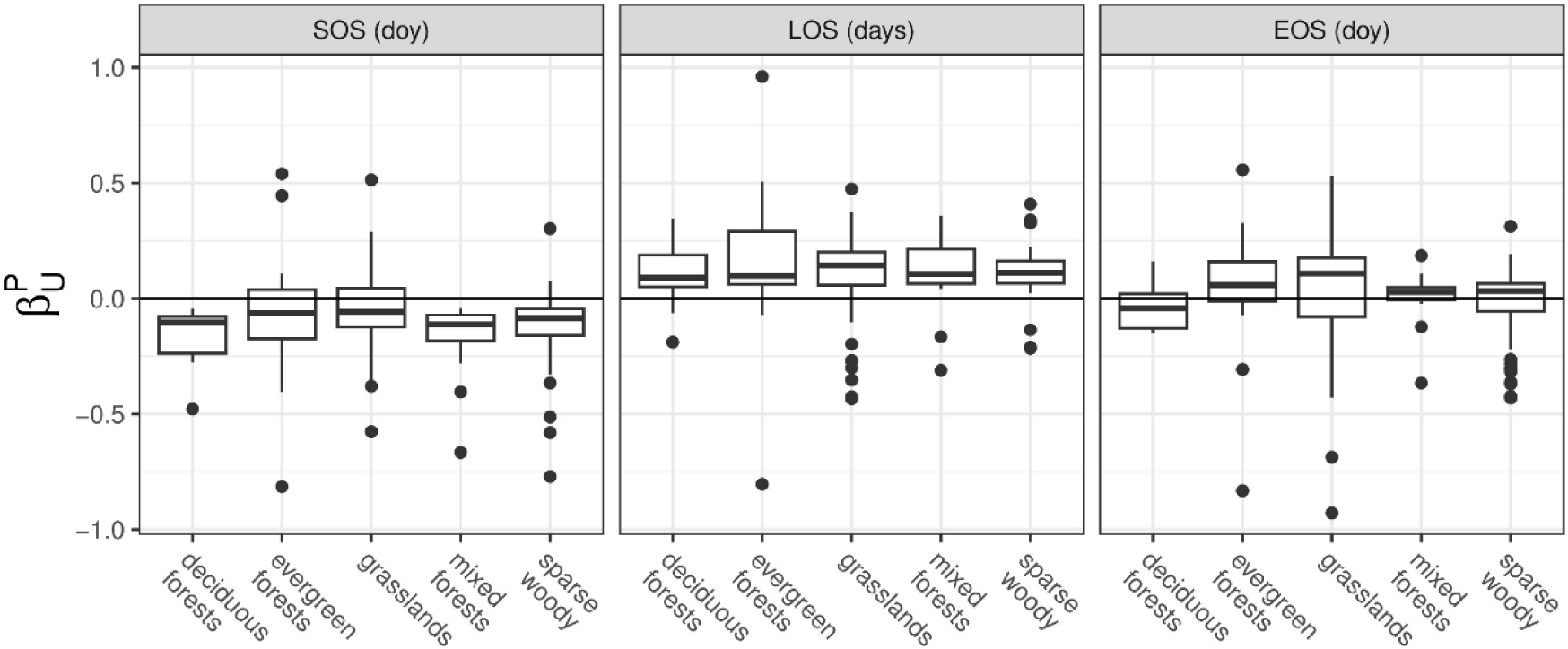
Phenological responses of the main vegetation types to urbanisation intensity expressed as the regression coefficient of the linear relationship between the phenological indexes and the proportion of urbanised area (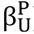). Year and elevation were included as covariables. We only show significant values, i.e., points represent regression coefficients that were significantly different from zero.

**Table 1.** Phenological response (mean and standard deviation) of the five vegetation types to urbanisation (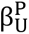) calculated using a multiple regression model including year and elevation as a covariable. We explored if the distribution of regression coefficients was significantly different from zero by running t-tests. SOS: start of season date, LOS: length of season in days, EOS: end of season date. df: degrees of freedom, p: p-value.

| | vegetation | $\beta_U^P$ | | | |
| --- | --- | --- | --- | --- | --- |
| | | mean ± sd | t | df | p |
| SOS<br>(doy) | evergreen forests | 0.02 ± 0.49 | 0.18 | 18 | 0.859 |
| | grasslands | -0.05 ± 0.18 | -1.89 | 45 | 0.065 |
| | mixed forests | -0.19 ± 0.23 | -4.36 | 28 | 0.000 |
| | sparse woody vegetation | -0.11 ± 0.14 | -6.38 | 63 | 0.000 |
| | deciduous forests | -0.16 ± 0.12 | -4.53 | 11 | 0.001 |
| LOS<br>(days) | evergreen forests | 0.15 ± 0.42 | 1.18 | 10 | 0.264 |
| | grasslands | 0.09 ± 0.2 | 3.39 | 56 | 0.001 |
| | mixed forests | 0.16 ± 0.27 | 3.47 | 30 | 0.002 |
| | sparse woody vegetation | 0.11 ± 0.11 | 7.27 | 49 | 0.000 |
| | deciduous forests | 0.1 ± 0.14 | 2.55 | 11 | 0.027 |
| EOS<br>(doy) | evergreen forests | 0.17 ± 0.64 | 1.03 | 13 | 0.322 |
| | grasslands | 0.05 ± 0.26 | 1.42 | 53 | 0.162 |
| | mixed forests | 0 ± 0.12 | 0.15 | 13 | 0.881 |
| | sparse woody vegetation | -0.02 ± 0.16 | -1.14 | 56 | 0.260 |
| | deciduous forests | -0.03 ± 0.1 | -0.96 | 11 | 0.357 |

The relationship between the end of season date (EOS) and urbanisation intensity resulted significantly positive in 41.80% of cases (i.e., delay of EOS) and significantly negative in 20.08% (i.e., advance in EOS) (Table 1; Figure 3). We found contrasting responses of EOS to urbanisation among vegetation types. The EOS of deciduous forests advanced on average 0.03 days per 1% increase in urbanisation intensity, whereas the EOS of evergreen forests, mixed forests and grasslands was delayed by 0.17, 0.01 and 0.05 days per unit increase in urbanisation intensity, respectively. The regression coefficients per metropolitan areas and vegetation type can be found in Appendix C, and the effect of year and elevation in Appendix D.

Annual mean temperature was a good explanatory variable of the strength of the response of SOS (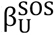) and EOS (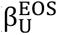) to urbanisation intensity in European metropolitan areas (Table 2; Figure 4). The advance of the SOS with increasing urbanisation was significantly most intense as annual mean temperature increased, whereas it tended to zero in colder metropolitan areas. We found a significant relationship between the effect of urbanisation on the EOS and annual mean temperature: urbanisation tended to delay the EOS of vegetation in metropolitan areas with annual mean temperatures below 11°C (i.e., metropolitan areas in the Temperate macrobioclimate), whereas it mostly brought forward the EOS of vegetation in metropolitan areas with annual mean temperatures over 11°C (i.e., metropolitan areas in the Mediterranean macrobioclimate). The uncertainty in the relationship between the phenological indexes and urbanisation intensity also increased with annual mean temperature (Figure 4), i.e., in metropolitan areas in the hottest regions there was a greater average phenological response to urbanisation intensity compared to colder areas, but also more variability associated to the model. In addition, minimum temperature of the coldest month and mean temperature of the coldest quarter were negatively correlated to 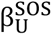 and 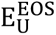 but are spatially positively correlated to annual mean temperature at a continental scale. 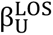 was negatively correlated to mean diurnal range, maximum temperature of the warmest month and mean temperature of the driest quarter. 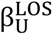 and 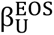 were positively correlated to precipitation of the driest month, driest quarter and warmest quarter, and negatively correlated to precipitation seasonality.

**Figure 4.**
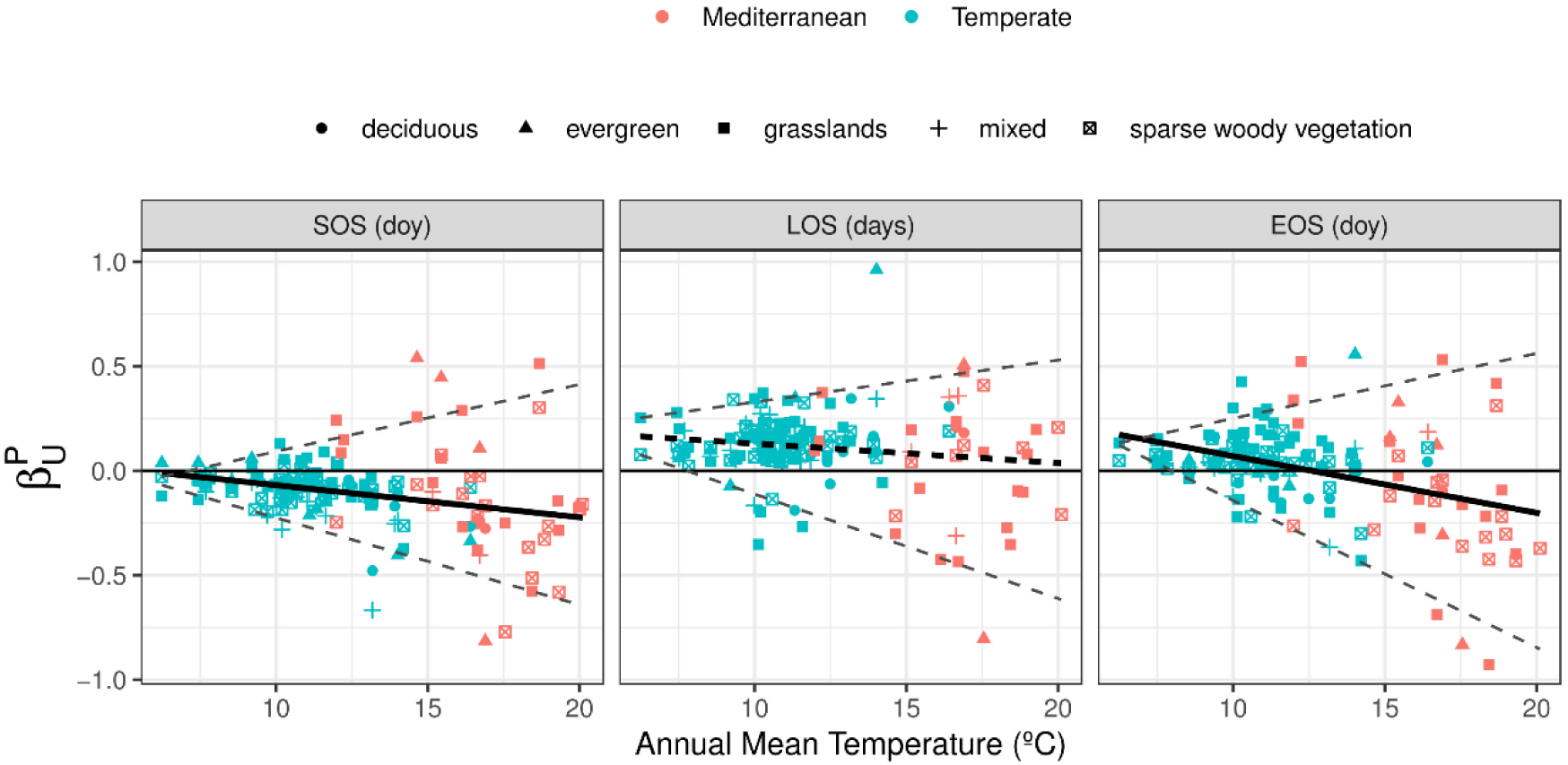
Variability of the phenological response of vegetation to urbanisation (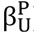) across an annual mean temperature gradient (mean annual mean temperature for the period 2002-2021). Colours indicate the macrobioclimate and symbols the main vegetation types. Bold solid lines represent significant linear regression patterns and dashed lines represent the 5 and 95 percentiles. SOS: start of season date, LOS: length of season in days, EOS: end of season date.

**Table 2.** Results of simple linear regressions using the phenological indexes as the response variable and each of the 19 BIOCLIM variables available from WorldClim as explanatory variables. BIOCLIM variables represent mean values for 2002-2021. SOS: start of season date, LOS: length of season in days, EOS: end of season date. P: p-value.

|  | SOS (doy) |  |  | LOS (days) |  |  | EOS (doy) |  |  |
| --- | --- | --- | --- | --- | --- | --- | --- | --- | --- |
| | $\beta_{bioclim}^P$ | R<sup>2</sup> | p | $\beta_{bioclim}^P$ | R<sup>2</sup> | p | $\beta_{bioclim}^P$ | R<sup>2</sup> | p |
| Annual Mean Temperature | -0.015 | 0.072 | 0.000 |  |  |  | -0.027 | 0.160 | 0.000 |
| Mean Diurnal Range |  |  |  | -0.022 | 0.021 | 0.038 |  |  |  |
| Isothermality |  |  |  |  |  |  | -0.013 | 0.071 | 0.001 |
| Temperature Seasonality | 0.000 | 0.061 | 0.001 |  |  |  | 0.000 | 0.033 | 0.015 |
| Max Temperature of Warmest Month |  |  |  | -0.009 | 0.020 | 0.041 | -0.016 | 0.065 | 0.001 |
| Min Temperature of Coldest Month | -0.015 | 0.110 | 0.000 |  |  |  | -0.020 | 0.144 | 0.000 |
| Temperature Annual Range | 0.012 | 0.065 | 0.001 |  |  |  |  |  |  |
| Mean Temperature of Wettest Quarter |  |  |  |  |  |  |  |  |  |
| Mean Temperature of Driest Quarter |  |  |  | -0.005 | 0.028 | 0.020 | -0.009 | 0.104 | 0.000 |
| Mean Temperature of Warmest Quarter | -0.010 | 0.026 | 0.020 |  |  |  | -0.022 | 0.100 | 0.000 |
| Mean Temperature of Coldest Quarter | -0.014 | 0.097 | 0.000 |  |  |  | -0.022 | 0.163 | 0.000 |
| Annual Precipitation | 0.000 | 0.032 | 0.011 | 0.000 | 0.023 | 0.032 |  |  |  |
| Precipitation of Wettest Month | -0.002 | 0.065 | 0.000 |  |  |  | -0.001 | 0.027 | 0.024 |
| Precipitation of Driest Month |  |  |  | 0.006 | 0.059 | 0.001 | 0.006 | 0.046 | 0.005 |
| Precipitation Seasonality |  |  |  | -0.004 | 0.083 | 0.000 | -0.005 | 0.109 | 0.000 |
| Precipitation of Wettest Quarter | -0.001 | 0.055 | 0.001 |  |  |  |  |  |  |
| Precipitation of Driest Quarter |  |  |  | 0.001 | 0.063 | 0.001 | 0.001 | 0.050 | 0.004 |
| Precipitation of Warmest Quarter |  |  |  | 0.001 | 0.047 | 0.003 | 0.001 | 0.092 | 0.000 |
| Precipitation of Coldest Quarter |  |  |  |  |  |  | 0.000 | 0.021 | 0.043 |

The phenological response of the different vegetation types to urbanisation (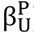) depended on the regional macrobioclimate (Table 3; Figure 5; results of random forest are available in Appendix E). In metropolitan areas from the Temperate macrobioclimate, we found consistent advances in the SOS, increased LOS and delays in the EOS across vegetation types with increasing urbanisation intensity (except for the advance in the EOS of deciduous forests). Vegetation from the Mediterranean macrobioclimate showed less consistent responses to urbanisation.

**Figure 5.**
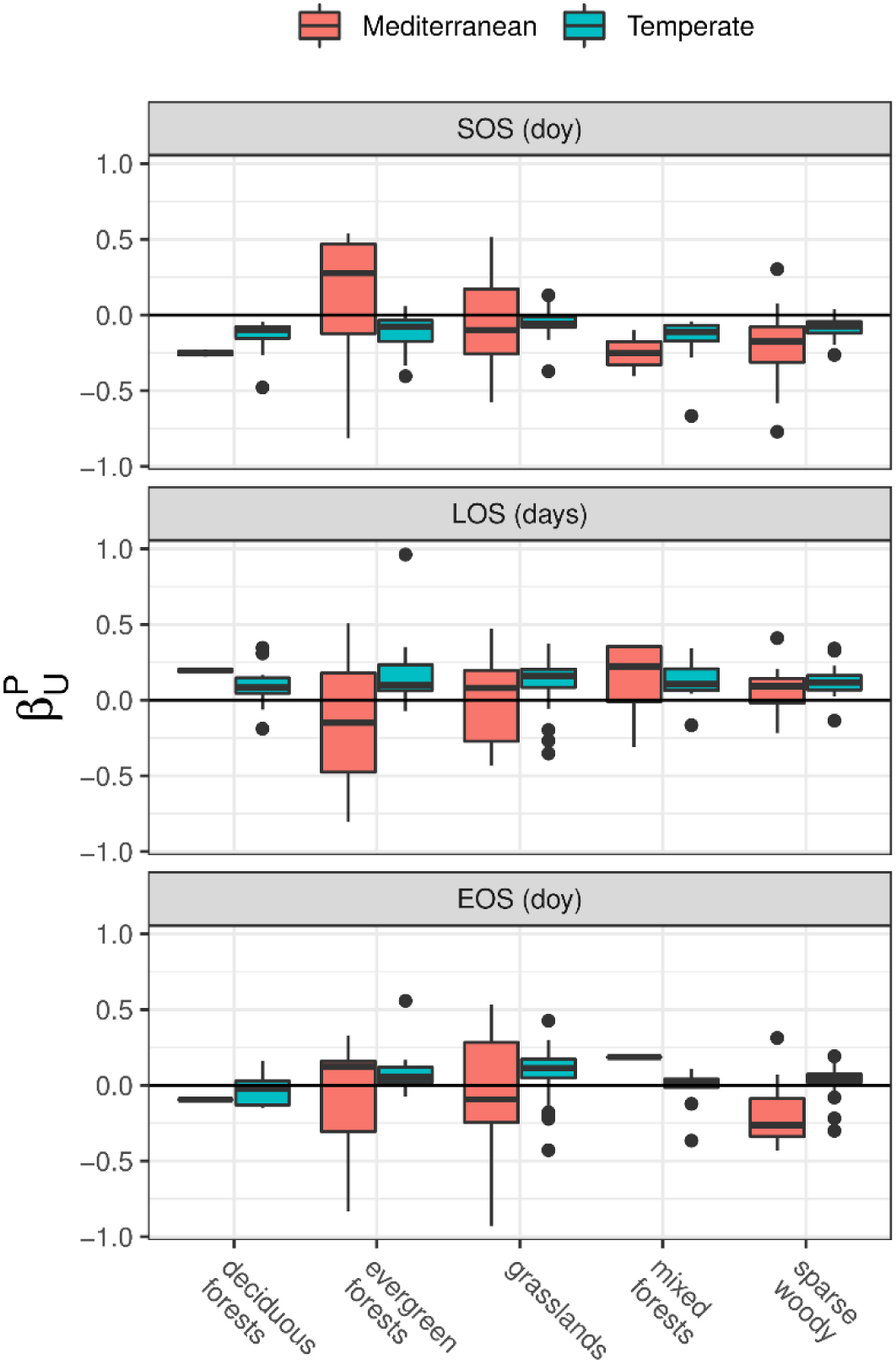
Phenological responses to urbanisation by the main vegetation types in metropolitan areas in the Temperate and Mediterranean macrobioclimates, expressed as the regression coefficient of the linear relationship between the phenological indexes and urbanisation intensity (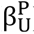). Year and elevation were included as covariables. We only show significant values, i.e., regression coefficients that were significantly different from zero.

**Table 3.**
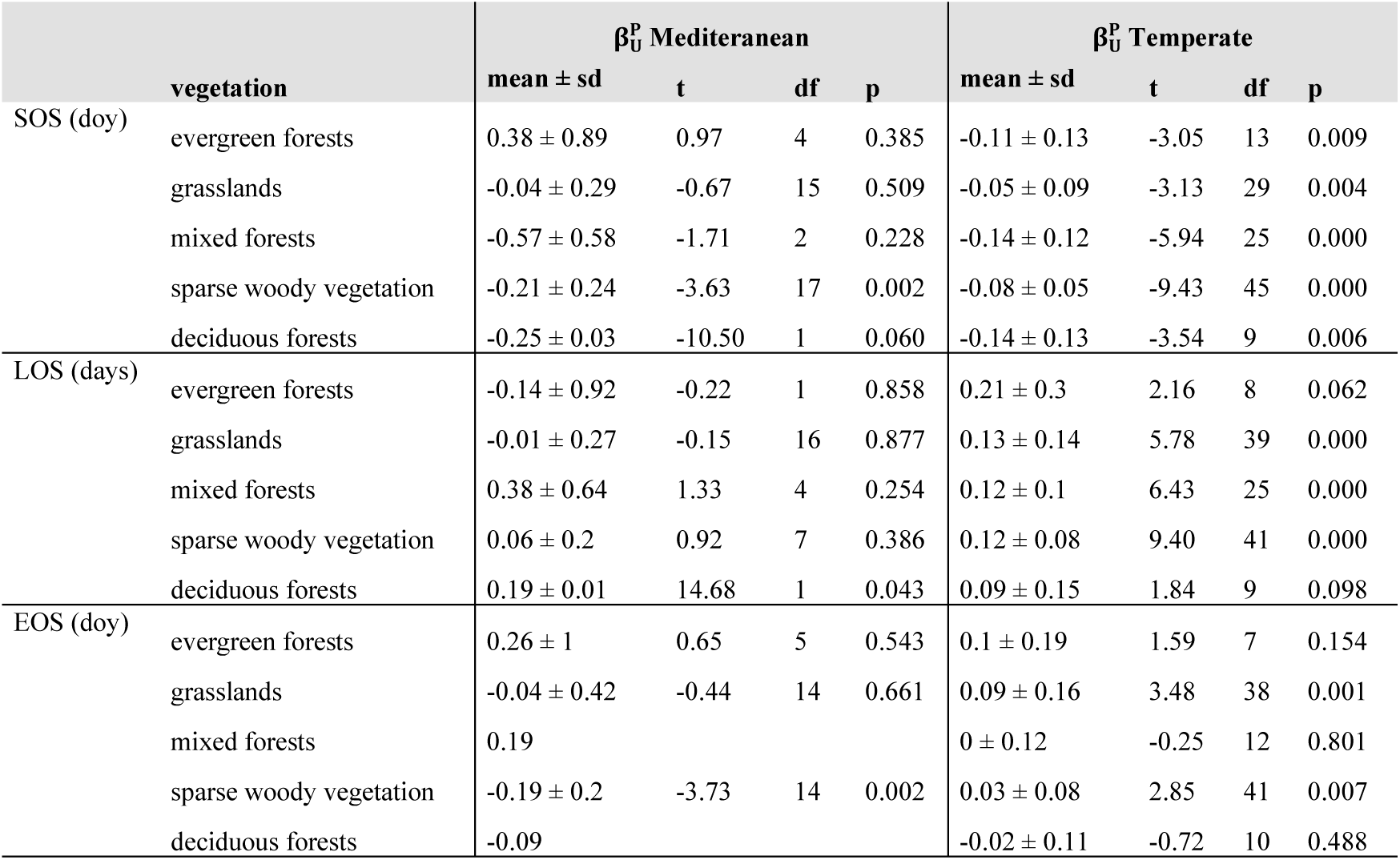
Phenological response (mean and standard deviation) to urbanisation and time of the main vegetation types in Temperate and Mediterranean metropolitan areas expressed as the regression coefficients (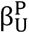 and 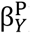) obtained from multiple regression models using elevation as a covariable. We explored whether the distribution of regression coefficients was significantly different from zero by running t-tests. There was only one observation of deciduous and mixed forests in metropolitan areas from the Mediterranean macrobioclimate. SOS: start of season date, LOS: length of season in days, EOS: end of season date. df: degrees of freedom, p: p-value.

The average winter and spring LST was negatively correlated with SOS across vegetation types (Figure 6): the SOS advanced 6.58 ± 6.42 days/K (mean and standard deviation) in vegetation from Mediterranean metropolitan areas, whereas in Temperate metropolitan areas the relationship was weaker but more homogeneous (-2.45 ± 1.49 days/K). The relationship between LST and urbanisation varied temporally (i.e., between day and night and along the year) and spatially (i.e., across Temperate and Mediterranean Europe; Appendix F).

**Figure 6.**
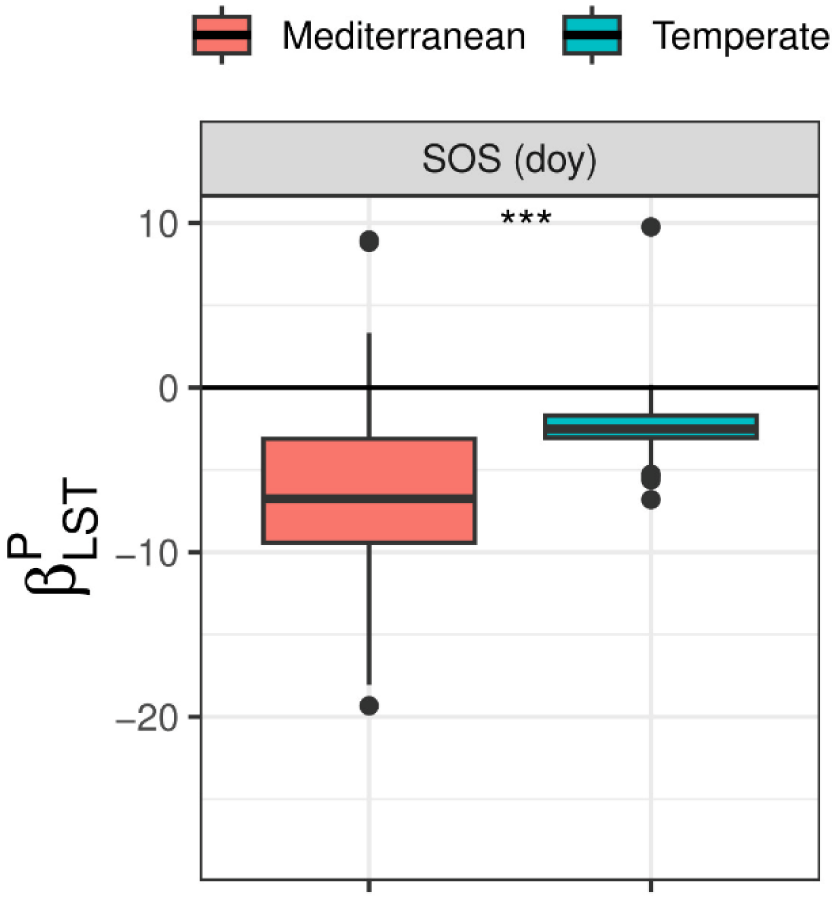
Response of the SOS to land surface temperature (LST) in the Temperate and Mediterranean macrobioclimates, expressed as the regression coefficient of the linear relationship between SOS and LST (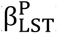). Year and elevation were included as covariables. We only show significant values, i.e., regression coefficients that were significantly different from zero. Asterisks denote results of t-test (*** p-value < 0.005).

## Discussion

### General response of vegetation phenology to urbanisation

Our results showed the general continuous pattern of phenological change in response to urbanisation at the landscape level within the largest European metropolitan areas and considering the entire Temperate-Mediterranean gradient. We report that the start of season date (SOS) of deciduous, evergreen and mixed forests, sparse woody vegetation and grasslands across metropolitan areas of Europe occurred 3-19 days earlier, and the LOS increased 10-16 days, in completely urbanised areas compared to the surrounding natural areas. These patterns are very consistent with those reported across regions by most studies (Li et al., 2019; Meng et al., 2020; Pan et al., 2021; Wohlfahrt et al., 2019; Yang et al., 2020). According to our analyses increasing preseason temperatures plays an important role, probably by breaking ecodormancy earlier (reviewed in Piao et al., 2023).

The end of season date (EOS) was frequently delayed (1-13 days) in completely urbanised areas. Yet, the relationship was less consistent than for SOS supporting the complex relationship between autumn phenophases and urbanisation (reviewed in Jochner & Menzel, 2015). This relationship was very dependent on the vegetation type. Most remarkably, the response of EOS of deciduous forests to urbanisation was mostly negative. It has been observed that the dates of leaf senescence have advanced in broadleaf trees from Central Europe and North America (Keenan and Richardson, 2015; Zani et al., 2020). A potential explanation is that earlier spring phenophases counteract the influence of warmer autumn temperatures via structural and physiological constraints on leaf longevity, or an increase probability of frost damage to buds and leaves (Keenan and Richardson, 2015). Earlier leaf-out has been demonstrated to lead to an advance in plants’ seasonal cycles via increased photosynthetic activity, causing earlier carbon saturation and inducing leaf senescence (Zani et al., 2020). This makes strongly inadvisable to group observations by wider ecosystem types as frequently occurs in urban phenological studies (e.g., Jia et al., 2021; Yang et al., 2020; Y. Zhang et al., 2022). As we discuss below, the response of EOS to urbanisation also depended on regional temperature which underlines the remarkable influence of macrobioclimatic factors.

### Spatial differences in the phenological response to urbanisation: effect of macrobioclimate

Our findings suggest that the SOS of deciduous, evergreen and mixed forests, sparse woody vegetation and grasslands across Europe advances with increasing urbanisation, although the strength and uncertainty of this relationship increased with annual mean temperature. The most marked SOS advances were registered in metropolitan areas from the Mediterranean region, but this is also where there was more uncertainty in the relationship between SOS and urbanisation. First, we found that the effect of urbanisation intensity on LST (related to the urban heat island) was stronger in metropolitan areas in the Temperate macrobioclimate throughout the day and the year, as noted in previous studies (Ma et al., 2016; Wienert and Kuttler, 2005). In Europe, we can therefore expect a more consistent and homogeneous phenological response to urbanisation in metropolitan areas at northern latitudes. In this regard, our results complement those of Li et al. (2019) who, using population density as a proxy for urbanisation, found a negative relationship between SOS and urbanisation across cold areas of Europe and North America. Second, this uncertainty might reflect physiological stress caused by increasing preseason and spring temperatures (Fu et al., 2015) and the interplay of other abiotic factors which may play an important role at lower latitudes such as water availability or proximity to the coast (Richardson et al., 2013; Yang et al., 2020). Finally, species turnover has been demonstrated to be a key factor to explain phenological differences in response to urbanisation (Alonzo et al., 2023). Therefore, differences in the diversity of urban and natural species pools might affect the relationship between vegetation phenology and urbanisation. In Mediterranean regions, where non-native species are widely planted within cities (Bayón et al., 2021), we might expect greater phenological differences and uncertainties between urban and natural landscapes. Meng et al. (2020) also argued that exotic species in urban ecosystems altered the phenological signals, but this aspect would not mask the general real pattern.

Regional annual mean temperature affected the strength, direction and uncertainty of the response of the EOS of European vegetation to urbanisation. The EOS of all vegetation types was positively correlated with urbanisation intensity in metropolitan areas with annual mean temperatures below 11°C (most metropolitan areas in the Temperate macroclimate), except for deciduous forest as indicated above. In metropolitan areas with annual mean temperatures over 11°C (most metropolitan areas in the Mediterranean macroclimate), the response was more heterogeneous but mostly negative, i.e., the EOS advanced in highly urbanised areas. Garonna et al. (2014) also found that, as opposed to vegetation in northern Europe, the EOS of vegetation in the Mediterranean microclimate advanced around 7.5 days during the period 1982-2011, also showing great spatial heterogeneity. Increasing summer and autumn temperatures and reduced water availability as a result of urbanisation might increase plant stress at the end of the growing season limiting plant growth, especially dry regions of the Northern Hemisphere where water is already a limiting factor (Menzel et al., 2020; Richardson et al., 2013), and consequently advancing the EOS (Liu et al., 2016). This is supported by our results, as the greatest delays of EOS were reported in metropolitan areas with the highest precipitation during summer.

### Uncertainties

Detection of phenophases using remote sensing is inevitably less precise than ground observations because it is affected by observation conditions, the choice of vegetation indexes and curve-fitting methods (Zeng et al., 2020). Li et al. (2019) and Wohlfahrt et al. (2019), using species-level data from the Pan European Phenology Database (PEP725), found that leaf-out occurred 1.0-4.3 days earlier in completely urbanised compared to non-urbanised areas and that leaf senescence occurred 1.3–2.7 days later. It has been demonstrated that highly heterogeneous pixels interfere in the phenological signal (Alonzo et al., 2023) and, in coarse resolution remote sensing data, SOS detection is biased towards earlier dates in areas with higher diversity of spring phenology dates per pixel (Tian et al., 2020; Zhang et al., 2017). Therefore, our results are probably overestimated due to the use of phenological data at a coarse resolution (i.e., 500 m). Also, MODIS phenological data (MCD12Q2 v006) is derived from the enhanced vegetation index (EVI2), which is known to show low seasonal variation in arid and evergreen ecosystems compared to other vegetation types (Jia et al., 2021). This might explain the less consistent pattern observed in Mediterranean evergreen forests and grasslands compared to other vegetation types.

### Conclusions

Remotely sensed data allowed us to detect phenological changes at the landscape level in response to urbanisation across the most populated European metropolitan areas. This contribution is among the first to report consistent responses of vegetation to urbanisation across metropolitan areas located along the entire Mediterranean-Temperate gradient and explore the macrobioclimatic factors that drive the observed spatial variability. It is worth noting that both the strength and direction of the effect of urbanisation on plant phenology varied at the continental scale and showed greater uncertainty in metropolitan areas in the Mediterranean macrobioclimate. Understanding the processes involved in the different responses of the diverse functional vegetation types requires applying process-based mechanistic models to define the phenological behaviour at the individual level. Our results suggest that macrobioclimatic constraints operating at the continental scale play an important role in determining the direction of the effect of urbanisation intensity on autumn phenophases. Future studies should focus on the ecological consequences of phenological shifts on urban ecosystems, such as changes in the pollen exposure of species with high allergenic potential and impacts on the populations of other organisms.

## Author’s contributions

JGD: Conceptualisation, Methodology, Formal analysis, Investigation, Data curation, Writing - original draft, Writing - review and editing. AMGB: Investigation, Writing - review and editing. JR: Conceptualisation, Methodology, Formal analysis, Investigation, Writing - review and editing.

## Data availability

This work was developed using open source datasets. The codes used in this study are available at GitHub (https://github.com/galanzse/europe_urbantrees).

## References

Alonzo, M., Baker, M.E., Caplan, J.S., Williams, A., Elmore, A.J., 2023. Canopy composition drives variability in urban growing season length more than the heat island effect. Sci. Total Environ. 884, 163818. 10.1016/j.scitotenv.2023.163818

Bayón, Á., Godoy, O., Maurel, N., van Kleunen, M., Vilà, M., 2021. Proportion of non-native plants in urban parks correlates with climate, socioeconomic factors and plant traits. Urban For. Urban Green. 63, 127215. 10.1016/j.ufug.2021.127215

Bernard-Verdier, M., Seitz, B., Buchholz, S., Kowarik, I., Lasunción Meijía, S., Jeschke, J.M., Lasunción Mejía, S., Jeschke, J.M., 2022. Grassland allergenicity increases with urbanisation and plant invasions. Ambio 2261–2277. 10.1007/s13280-022-01741-z

Booth, T.H., Nix, H.A., Busby, J.R., Hutchinson, M.F., 2014. bioclim: the first species distribution modelling package, its early applications and relevance to most current MaxEnt studies. Divers. Distrib. 20, 1–9. 10.1111/ddi.12144

Czaja, M., Kołton, A., Muras, P., 2020. The complex issue of urban trees-stress factor accumulation and ecological service possibilities. Forests 11, 1–24. 10.3390/F11090932

Fick, S.E., Hijmans, R.J.R.J., 2017. WorldClim 2: new 1km spatial resolution climate surfaces for global land areas. Int. J. Climatol. 37, 4302–4315. 10.1002/joc.5086

Fu, Y.H., Zhao, H., Piao, S., Peaucelle, M., Peng, S., Zhou, G., Ciais, P., Huang, M., Menzel, A., Peñuelas, J., Song, Y., Vitasse, Y., Zeng, Z., Janssens, I.A., 2015. Declining global warming effects on the phenology of spring leaf unfolding. Nature 526, 104–107. 10.1038/nature15402

Galán Díaz, J., Gutierrez-Bustillo, A.M., Rojo, J., 2023. Influence of urbanisation on the phenology of evergreen coniferous and deciduous broadleaf trees in Madrid (Spain). Landsc. Urban Plan. 235. 10.1016/j.landurbplan.2023.104760

Garcia, R.A., Cabeza, M., Rahbek, C., Araújo, M.B., 2014. Multiple Dimensions of Climate Change and Their Implications for Biodiversity. Science (80-.). 344. 10.1126/science.1247579

Garonna, I., de Jong, R., de Wit, A.J.W., Mücher, C.A., Schmid, B., Schaepman, M.E., 2014. Strong contribution of autumn phenology to changes in satellite-derived growing season length estimates across Europe (1982-2011). Glob. Chang. Biol. 20, 3457–3470. 10.1111/gcb.12625

Greenwell, Brandon, M., 2017. pdp: An R Package for Constructing Partial Dependence Plots. R J. 9, 421. 10.32614/RJ-2017-016

Hao, L., Huang, X., Qin, M., Liu, Y., Li, W., Sun, G., 2018. Ecohydrological Processes Explain Urban Dry Island Effects in a Wet Region, Southern China. Water Resour. Res. 54, 6757– 6771. 10.1029/2018WR023002

Harris, I., Osborn, T.J., Jones, P., Lister, D., 2020. Version 4 of the CRU TS monthly high-resolution gridded multivariate climate dataset. Sci. Data 7, 109. 10.1038/s41597-020-0453-3

Hijmans, R., Phillips, S., Leathwick, J., Elith, J., 2023. dismo: Species Distribution Modeling. R package version 1.3–14.

Hijmans, R.J., Bivand, R., Forner, K., Ooms, J., Pebesma, E., Sumner, M.D., 2022. Package “terra.”

Hufkens, K., 2022. The MODISTools package: an interface to the MODIS Land Products Subsets Web Services.

Inouye, D.W., 2022. Climate change and phenology. Wiley Interdiscip. Rev. Clim. Chang. 13, 1–17. 10.1002/wcc.764

Jia, W., Zhao, S., Zhang, X., Liu, S., Henebry, G.M., Liu, L., 2021. Urbanization imprint on land surface phenology: The urban–rural gradient analysis for Chinese cities. Glob. Chang. Biol. 27, 2895–2904. 10.1111/gcb.15602

Jochner, S., Menzel, A., 2015. Urban phenological studies – Past, present, future. Environ. Pollut. 203, 250–261. 10.1016/j.envpol.2015.01.003

Kabano, P., Lindley, S., Harris, A., 2021. Evidence of urban heat island impacts on the vegetation growing season length in a tropical city. Landsc. Urban Plan. 206, 103989. 10.1016/j.landurbplan.2020.103989

Keenan, T.F., Richardson, A.D., 2015. The timing of autumn senescence is affected by the timing of spring phenology: implications for predictive models. Glob. Chang. Biol. 21, 2634–2641. 10.1111/gcb.12890

Li, D., Stucky, B.J., Deck, J., Baiser, B., Guralnick, R.P., 2019. The effect of urbanization on plant phenology depends on regional temperature. Nat. Ecol. Evol. 3, 1661–1667. 10.1038/s41559-019-1004-1

Liu, Q., Fu, Y.H., Zhu, Z., Liu, Y., Liu, Z., Huang, M., Janssens, I.A., Piao, S., 2016. Delayed autumn phenology in the Northern Hemisphere is related to change in both climate and spring phenology. Glob. Chang. Biol. 22, 3702–3711. 10.1111/gcb.13311

Ma, Q., Wu, J., He, C., 2016. A hierarchical analysis of the relationship between urban impervious surfaces and land surface temperatures: spatial scale dependence, temporal variations, and bioclimatic modulation. Landsc. Ecol. 31, 1139–1153. 10.1007/s10980-016-0356-z

Meng, L., Mao, J., Zhou, Y., Richardson, A.D., Lee, X., Thornton, P.E., Ricciuto, D.M., Li, X., Dai, Y., Shi, X., Jia, G., 2020. Urban warming advances spring phenology but reduces the response of phenology to temperature in the conterminous United States. Proc. Natl. Acad. Sci. U. S. A. 117, 4228–4233. 10.1073/pnas.1911117117

Menzel, A., Yuan, Y., Matiu, M., Sparks, T., Scheifinger, H., Gehrig, R., Estrella, N., 2020. Climate change fingerprints in recent European plant phenology. Glob. Chang. Biol. 26, 2599–2612. 10.1111/gcb.15000

Neil, K., Wu, J., 2006. Effects of urbanization on plant flowering phenology: A review. Urban Ecosyst. 9, 243–257. 10.1007/s11252-006-9354-2

Pan, L., Xia, H., Yang, J., Niu, W., Wang, R., Song, H., Guo, Y., Qin, Y., 2021. Mapping cropping intensity in Huaihe basin using phenology algorithm, all Sentinel-2 and Landsat images in Google Earth Engine. Int. J. Appl. Earth Obs. Geoinf. 102, 102376. 10.1016/j.jag.2021.102376

Piao, S., Liu, Q., Chen, A., Janssens, I.A., Fu, Y., Dai, J., Liu, L., Lian, X., Shen, M., Zhu, X., 2023. Plant phenology and global climate change: Current progresses and challenges. Glob. Chang. Biol. 25, 161109. 10.1016/j.scitotenv.2022.161109

Ren, Q., He, C., Huang, Q., Zhou, Y., 2018. Urbanization impacts on vegetation phenology in China. Remote Sens. 10, 1–16. 10.3390/rs10121905

Renner, S.S., Zohner, C.M., 2018. Climate change and phenological mismatch in trophic interactions among plants, insects, and vertebrates. Annu. Rev. Ecol. Evol. Syst. 49, 165–182. 10.1146/annurev-ecolsys-110617-062535

Richardson, A.D., Keenan, T.F., Migliavacca, M., Ryu, Y., Sonnentag, O., Toomey, M., 2013. Climate change, phenology, and phenological control of vegetation feedbacks to the climate system. Agric. For. Meteorol. 169, 156–173. 10.1016/j.agrformet.2012.09.012

Rivas-Martínez, S., Rivas-Saenz, S., Penas, A., 2002. Worldwide bioclimatic classification system. Backhuys Pub., Kerkwerve, The Netherlands.

Rojo, J., Oteros, J., Picornell, A., Maya-Manzano, J.M., Damialis, A., Zink, K., Werchan, M., Werchan, B., Smith, M., Menzel, A., Timpf, S., Traidl-Hoffmann, C., Bergmann, K.C., Schmidt-Weber, C.B., Buters, J., 2021. Effects of future climate change on birch abundance and their pollen load. Glob. Chang. Biol. 27, 5934–5949. 10.1111/gcb.15824

Sismanidis, P., Bechtel, B., Perry, M., Ghent, D., 2022. The Seasonality of Surface Urban Heat Islands across Climates. Remote Sens. 14, 1–21. 10.3390/rs14102318

Templ, B., Koch, E., Bolmgren, K., Ungersböck, M., Paul, A., Scheifinger, H., Rutishauser, T., Busto, M., Chmielewski, F.-M., Hájková, L., Hodzić, S., Kaspar, F., Pietragalla, B., Romero-Fresneda, R., Tolvanen, A., Vučetič, V., Zimmermann, K., Zust, A., 2018. Pan European Phenological database (PEP725): a single point of access for European data. Int. J. Biometeorol. 62, 1109–1113. 10.1007/s00484-018-1512-8

Tian, J., Zhu, X., Wu, J., Shen, M., Chen, J., 2020. Coarse-resolution satellite images overestimate urbanization effects on vegetation spring phenology. Remote Sens. 12. 10.3390/RS12010117

Trusilova, K., Churkina, G., 2008. The response of the terrestrial biosphere to urbanization: land cover conversion, climate, and urban pollution. Biogeosciences 5, 1505–1515. 10.5194/bg-5-1505-2008

Wienert, U., Kuttler, W., 2005. The dependence of the urban heat island intensity on latitude - A statistical approach. Meteorol. Zeitschrift 14, 677–686. 10.1127/0941-2948/2005/0069

Wohlfahrt, G., Tomelleri, E., Hammerle, A., 2019. The urban imprint on plant phenology. Nat. Ecol. Evol. 3, 1668–1674. 10.1038/s41559-019-1017-9

Wu, C., Wang, X., Wang, H., Ciais, P., Peñuelas, J., Myneni, R.B., Desai, A.R., Gough, C.M., Gonsamo, A., Black, A.T., Jassal, R.S., Ju, W., Yuan, W., Fu, Y., Shen, M., Li, S., Liu, R., Chen, J.M., Ge, Q., 2018. Contrasting responses of autumn-leaf senescence to daytime and night-time warming. Nat. Clim. Chang. 8, 1092–1096. 10.1038/s41558-018-0346-z

Yang, J., Luo, X., Jin, C., Xiao, X., Xia, J., 2020. Spatiotemporal patterns of vegetation phenology along the urban–rural gradient in Coastal Dalian, China. Urban For. Urban Green. 54, 126784. 10.1016/j.ufug.2020.126784

Yu, H., Luedeling, E., Xu, J., 2010. Winter and spring warming result in delayed spring phenology on the Tibetan Plateau. Proc. Natl. Acad. Sci. 107, 22151–22156. 10.1073/pnas.1012490107

Zani, D., Crowther, T.W., Mo, L., Renner, S.S., Zohner, C.M., 2020. Increased growing-season productivity drives earlier autumn leaf senescence in temperate trees. Science (80-.). 370, 1066–1071. 10.1126/science.abd8911

Zeng, L., Wardlow, B.D., Xiang, D., Hu, S., Li, D., 2020. A review of vegetation phenological metrics extraction using time-series, multispectral satellite data. Remote Sens. Environ. 237, 111511. 10.1016/j.rse.2019.111511

Zhang, X., Wang, J., Gao, F., Liu, Y., Schaaf, C., Friedl, M., Yu, Y., Jayavelu, S., Gray, J., Liu, L., Yan, D., Henebry, G.M., 2017. Exploration of scaling effects on coarse resolution land surface phenology. Remote Sens. Environ. 190, 318–330. 10.1016/j.rse.2017.01.001

Zhang, Y., Yin, P., Li, X., Niu, Q., Wang, Y., Cao, W., Huang, J., Chen, H., Yao, X., Yu, L., Li, B., 2022. The divergent response of vegetation phenology to urbanization: A case study of Beijing city, China. Sci. Total Environ. 803, 150079. 10.1016/j.scitotenv.2021.150079

Zhao, M., Cheng, C., Zhou, Y., Li, X., Shen, S., Song, C., 2022. A global dataset of annual urban extents (1992-2020) from harmonized nighttime lights. Earth Syst. Sci. Data 14, 517–534. 10.5194/essd-14-517-2022

Ziello, C., Estrella, N., Kostova, M., Koch, E., Menzel, A., 2009. Influence of altitude on phenology of selected plant species in the Alpine region (1971–2000). Clim. Res. 39, 227–234. 10.3354/cr00822

