## Supplementary material for "The phenological response of European vegetation to urbanisation is mediated by macrobioclimatic factors"

**Appendix A.** Effect of different sources of urbanisation intensity (CORINE, imperviousness) and buffer sizes (250m, 500m, 1000m) on the start of season date (SOS). We run multiple regression models using SOS and urbanisation intensity as the response and explanatory variables respectively. Elevation and year were included as covariables. We found a high linear correlation between the regression coefficients ($\beta_U^{SOS}$). We used CORINE Land Cover at 500 m for subsequent analyses.

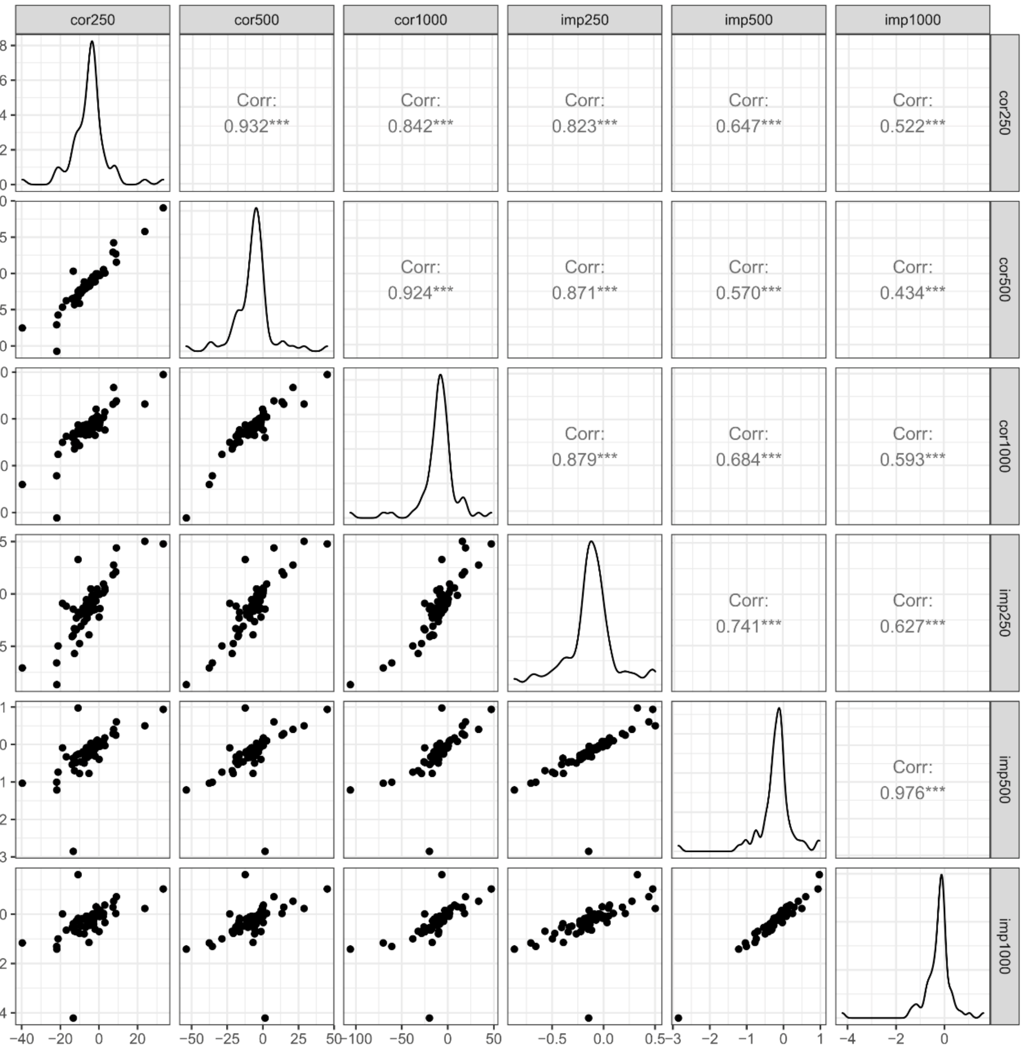

**Appendix B.** Average start of season date (SOS), length of season (LOS) and end of season date (EOS) of the five vegetation types in Europe considered in this study.

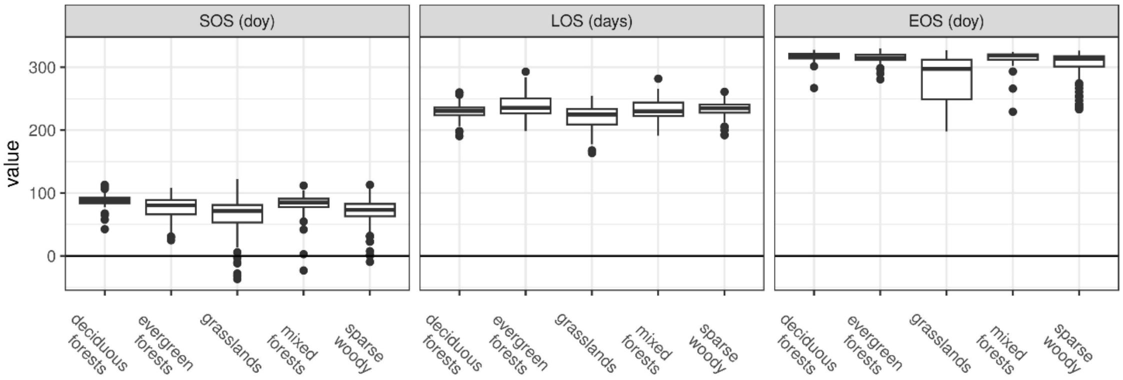

**Appendix C.** Results of multiple linear regression with phenological indexes as response variables and urbanisation intensity, year and elevation as explanatory variables. Urbanisation intensity was calculated as the proportion of built-up area extracted from CORINE using a 500 m circular buffer. We show regression coefficients of significant relationships between the phenological indexes and urbanisation ($\beta_U^P$) (i.e., $p < 0.05$). "n.s." indicates a non-significant relationship; empty cells indicate insufficient sample size (i.e., less than 50 observations). SOS: start of season date, LOS: length of season, EOS: end of season date. Deci: deciduous forests, ever: evergreen forests, spar: sparse woody vegetation, gras: grasslands. mixe: mixed forests,

| | | | | | $\beta_U^P$ | | |
|---|---|---|---|---|---|---|---|
| **name** | **macroclim** | **pheno** | **deci** | **ever** | **spar** | **gras** | **mixe** |
| Adana | Mediterranean | SOS | | -0.17 | | -0.18 | |
| Ankara (MA) | Mediterranean | SOS | | 0.15 | | n.s. | |
| Antalya | Mediterranean | SOS | | n.s. | | -0.26 | |
| Athens | Mediterranean | SOS | n.s. | n.s. | | -0.33 | |
| Barcelona | Mediterranean | SOS | 0.11 | -0.25 | -0.40 | -0.02 | |
| Belgrade | Temperate | SOS | | -0.16 | | -0.14 | n.s. |
| Berlin | Temperate | SOS | -0.07 | 0.05 | -0.07 | 0.01 | n.s. |
| Birmingham | Temperate | SOS | | n.s. | -0.28 | -0.19 | |
| Bremen | Temperate | SOS | | 0.13 | -0.10 | -0.04 | |
| Brussels | Temperate | SOS | -0.08 | -0.05 | -0.07 | -0.11 | n.s. |
| Bucharest | Temperate | SOS | | 0.03 | | 0.04 | n.s. |
| Budapest | Temperate | SOS | | n.s. | | -0.09 | -0.04 |
| Bursa | Mediterranean | SOS | 0.54 | 0.26 | | -0.07 | |
| Copenhagen | Temperate | SOS | | -0.07 | -0.21 | -0.20 | n.s. |
| Dresden | Temperate | SOS | -0.21 | n.s. | -0.10 | -0.05 | |
| Dublin | Temperate | SOS | n.s. | n.s. | n.s. | -0.15 | |
| Edinburgh | Temperate | SOS | | n.s. | n.s. | -0.19 | |
| Eskişehir | Mediterranean | SOS | | 0.24 | | -0.25 | |
| Frankfurt | Temperate | SOS | n.s. | n.s. | -0.04 | -0.07 | n.s. |
| Gaziantep | Mediterranean | SOS | | 0.29 | | | |
| Genoa | Temperate | SOS | -0.34 | | n.s. | -0.08 | -0.27 |
| Glasgow | Temperate | SOS | n.s. | n.s. | n.s. | -0.13 | |
| Göteborg | Temperate | SOS | -0.05 | -0.08 | -0.05 | -0.10 | |
| Hamburg | Temperate | SOS | n.s. | n.s. | -0.08 | -0.04 | |
| Hannover | Temperate | SOS | n.s. | -0.06 | -0.13 | -0.11 | |
| Helsinki | Temperate | SOS | 0.04 | -0.12 | n.s. | -0.03 | |
| Istanbul (MA) | Mediterranean | SOS | n.s. | -0.06 | -0.10 | -0.16 | n.s. |
| İzmir | Mediterranean | SOS | n.s. | n.s. | | -0.37 | |
| Konya | Mediterranean | SOS | | 0.09 | | | |
| Kraków | Temperate | SOS | | -0.08 | -0.13 | -0.08 | -0.09 |
| Leipzig (MA) | Temperate | SOS | | -0.06 | n.s. | -0.08 | n.s. |
| Lisbon | Mediterranean | SOS | n.s. | -0.25 | | -0.77 | |
| Liverpool | Temperate | SOS | | n.s. | n.s. | -0.12 | |
| Łódź | Temperate | SOS | 0.06 | 0.01 | -0.06 | n.s. | n.s. |
| London | Temperate | SOS | -0.15 | -0.10 | -0.21 | -0.13 | n.s. |
| Lyon | Temperate | SOS | | -0.11 | -0.67 | -0.09 | -0.48 |
| Madrid | Mediterranean | SOS | 0.45 | 0.06 | | 0.08 | |
| Málaga | Mediterranean | SOS | | 0.51 | | 0.30 | |
| Marseille | Mediterranean | SOS | n.s. | n.s. | n.s. | -0.03 | |
| Mersin | Mediterranean | SOS | | n.s. | | -0.16 | |

| | | | | | | | |
|---|---|---|---|---|---|---|---|
| Milan | Temperate | SOS | -0.40 | n.s. | -0.24 | -0.05 | -0.11 |
| Munich | Temperate | SOS | | n.s. | n.s. | -0.05 | |
| Naples | Mediterranean | SOS | -0.81 | n.s. | -1.22 | -0.17 | -0.28 |
| Nürnberg | Temperate | SOS | -0.13 | 0.05 | -0.06 | 0.03 | |
| Oslo | Temperate | SOS | 0.04 | -0.14 | n.s. | -0.04 | |
| Palermo | Mediterranean | SOS | 1.67 | -0.14 | | n.s. | |
| Paris | Temperate | SOS | | -0.06 | -0.12 | -0.12 | -0.08 |
| Poznań | Temperate | SOS | n.s. | n.s. | n.s. | -0.07 | |
| Prague | Temperate | SOS | n.s. | -0.06 | n.s. | -0.06 | n.s. |
| Randstad (MA) | Temperate | SOS | | 0.06 | -0.15 | -0.07 | |
| Riga | Temperate | SOS | n.s. | -0.06 | -0.10 | -0.06 | |
| Rin-Ruhr (MA) | Temperate | SOS | n.s. | n.s. | -0.06 | -0.01 | n.s. |
| Rome | Mediterranean | SOS | n.s. | -0.38 | n.s. | -0.22 | -0.23 |
| Sarajevo | Temperate | SOS | | 0.09 | | -0.11 | -0.10 |
| Sevilla | Mediterranean | SOS | | -0.29 | | -0.58 | |
| Sheffield | Temperate | SOS | | n.s. | n.s. | -0.07 | n.s. |
| Skopje | Temperate | SOS | | -0.03 | | -0.16 | |
| Sofia | Temperate | SOS | | -0.01 | -0.17 | -0.05 | n.s. |
| Stockholm | Temperate | SOS | -0.03 | -0.05 | n.s. | -0.04 | |
| Stuttgart | Temperate | SOS | | n.s. | -0.11 | -0.07 | n.s. |
| Toulouse | Temperate | SOS | | -0.37 | | -0.26 | |
| Turin | Temperate | SOS | | n.s. | -0.25 | -0.07 | -0.17 |
| Valencia | Mediterranean | SOS | | -0.58 | | -0.51 | |
| Vienna | Temperate | SOS | -0.18 | -0.08 | -0.17 | -0.04 | -0.08 |
| Vilnius | Temperate | SOS | n.s. | -0.04 | n.s. | -0.01 | |
| Warsaw | Temperate | SOS | -0.05 | -0.02 | -0.04 | -0.04 | n.s. |
| Wrocław | Temperate | SOS | | -0.06 | -0.17 | -0.14 | n.s. |
| Zagreb | Temperate | SOS | | -0.13 | | -0.09 | -0.07 |
| Zaragoza | Mediterranean | SOS | | -0.27 | | -0.11 | |
| Adana | Mediterranean | LOS | | n.s. | | 0.21 | |
| Ankara (MA) | Mediterranean | LOS | | 0.37 | | n.s. | |
| Antalya | Mediterranean | LOS | | 0.08 | | n.s. | |
| Athens | Mediterranean | LOS | n.s. | -0.10 | | 0.11 | |
| Barcelona | Mediterranean | LOS | n.s. | -0.43 | 0.36 | n.s. | |
| Belgrade | Temperate | LOS | | 0.14 | | 0.19 | n.s. |
| Berlin | Temperate | LOS | 0.06 | 0.23 | 0.06 | 0.04 | n.s. |
| Birmingham | Temperate | LOS | | -0.20 | 0.27 | 0.33 | |
| Bremen | Temperate | LOS | | -0.35 | n.s. | n.s. | |
| Brussels | Temperate | LOS | n.s. | 0.19 | 0.05 | 0.11 | 0.08 |
| Bucharest | Temperate | LOS | | 0.14 | | n.s. | n.s. |
| Budapest | Temperate | LOS | | 0.07 | | 0.12 | 0.04 |
| Bursa | Mediterranean | LOS | n.s. | -0.30 | | -0.22 | |
| Copenhagen | Temperate | LOS | | 0.16 | 0.22 | 0.21 | n.s. |
| Dresden | Temperate | LOS | n.s. | 0.34 | 0.11 | 0.14 | |
| Dublin | Temperate | LOS | n.s. | n.s. | 0.27 | 0.22 | |
| Edinburgh | Temperate | LOS | | n.s. | n.s. | 0.34 | |
| Eskişehir | Mediterranean | LOS | | 0.10 | | n.s. | |
| Frankfurt | Temperate | LOS | n.s. | 0.13 | 0.05 | 0.07 | n.s. |
| Gaziantep | Mediterranean | LOS | | -0.43 | | | |
| Genoa | Temperate | LOS | n.s. | | n.s. | 0.19 | 0.31 |
| Glasgow | Temperate | LOS | n.s. | 0.07 | n.s. | 0.16 | |
| Göteborg | Temperate | LOS | 0.10 | 0.05 | 0.07 | 0.11 | |
| Hamburg | Temperate | LOS | n.s. | n.s. | 0.08 | 0.05 | |
| Hannover | Temperate | LOS | n.s. | 0.16 | 0.14 | 0.16 | |
| Helsinki | Temperate | LOS | n.s. | 0.25 | n.s. | 0.08 | |
| Istanbul (MA) | Mediterranean | LOS | n.s. | 0.20 | 0.09 | 0.04 | n.s. |
| İzmir | Mediterranean | LOS | n.s. | -0.27 | | n.s. | |
| Konya | Mediterranean | LOS | | 0.14 | | | |
| Kraków | Temperate | LOS | | 0.17 | 0.11 | 0.16 | n.s. |
| Leipzig (MA) | Temperate | LOS | | 0.08 | n.s. | -0.14 | n.s. |

| | | | | | | | |
|---|---|---|---|---|---|---|---|
| Lisbon | Mediterranean | LOS | -0.80 | 0.09 | | 0.41 | |
| Liverpool | Temperate | LOS | | 0.16 | n.s. | 0.23 | |
| Łódź | Temperate | LOS | -0.07 | 0.16 | 0.10 | 0.05 | n.s. |
| London | Temperate | LOS | 0.19 | 0.17 | 0.21 | 0.33 | n.s. |
| Lyon | Temperate | LOS | | n.s. | n.s. | n.s. | 0.35 |
| Madrid | Mediterranean | LOS | n.s. | -0.08 | | n.s. | |
| Málaga | Mediterranean | LOS | | -0.10 | | n.s. | |
| Marseille | Mediterranean | LOS | n.s. | n.s. | 0.35 | n.s. | |
| Mersin | Mediterranean | LOS | | n.s. | | -0.21 | |
| Milan | Temperate | LOS | 0.96 | n.s. | 0.34 | 0.06 | 0.09 |
| Munich | Temperate | LOS | | 0.17 | n.s. | 0.08 | |
| Naples | Mediterranean | LOS | 0.51 | 0.47 | 1.44 | 0.12 | 0.18 |
| Nürnberg | Temperate | LOS | 0.23 | 0.37 | 0.09 | 0.10 | |
| Oslo | Temperate | LOS | n.s. | 0.28 | n.s. | 0.12 | |
| Palermo | Mediterranean | LOS | n.s. | 0.20 | | n.s. | |
| Paris | Temperate | LOS | | 0.21 | 0.13 | 0.14 | 0.09 |
| Poznań | Temperate | LOS | n.s. | 0.16 | n.s. | 0.07 | |
| Prague | Temperate | LOS | n.s. | 0.23 | n.s. | 0.11 | n.s. |
| Randstad (MA) | Temperate | LOS | | 0.20 | 0.14 | 0.11 | |
| Riga | Temperate | LOS | n.s. | 0.08 | 0.19 | 0.09 | |
| Rin-Ruhr (MA) | Temperate | LOS | 0.09 | 0.18 | 0.04 | 0.04 | n.s. |
| Rome | Mediterranean | LOS | n.s. | 0.24 | -0.31 | 0.07 | 0.21 |
| Sarajevo | Temperate | LOS | | -0.27 | | 0.15 | n.s. |
| Sevilla | Mediterranean | LOS | | n.s. | | n.s. | |
| Sheffield | Temperate | LOS | | 0.34 | -0.17 | 0.19 | n.s. |
| Skopje | Temperate | LOS | | 0.14 | | n.s. | |
| Sofia | Temperate | LOS | | 0.24 | 0.17 | 0.13 | -0.19 |
| Stockholm | Temperate | LOS | 0.06 | 0.20 | 0.06 | 0.08 | |
| Stuttgart | Temperate | LOS | | 0.05 | 0.09 | 0.07 | n.s. |
| Toulouse | Temperate | LOS | | -0.06 | | n.s. | |
| Turin | Temperate | LOS | | 0.09 | n.s. | 0.13 | 0.16 |
| Valencia | Mediterranean | LOS | | -0.35 | | n.s. | |
| Vienna | Temperate | LOS | 0.35 | n.s. | 0.22 | 0.06 | 0.05 |
| Vilnius | Temperate | LOS | n.s. | 0.11 | n.s. | 0.02 | |
| Warsaw | Temperate | LOS | n.s. | 0.06 | 0.05 | 0.06 | n.s. |
| Wrocław | Temperate | LOS | | 0.17 | 0.22 | 0.17 | n.s. |
| Zagreb | Temperate | LOS | | 0.32 | | 0.16 | -0.06 |
| Zaragoza | Mediterranean | LOS | | n.s. | | n.s. | |
| Adana | Mediterranean | EOS | | n.s. | | n.s. | |
| Ankara (MA) | Mediterranean | EOS | | 0.52 | | n.s. | |
| Antalya | Mediterranean | EOS | | n.s. | | -0.30 | |
| Athens | Mediterranean | EOS | n.s. | -0.09 | | -0.22 | |
| Barcelona | Mediterranean | EOS | 0.12 | -0.69 | n.s. | -0.06 | |
| Belgrade | Temperate | EOS | | n.s. | | 0.04 | n.s. |
| Berlin | Temperate | EOS | -0.01 | 0.28 | n.s. | 0.06 | n.s. |
| Birmingham | Temperate | EOS | | -0.14 | n.s. | 0.14 | |
| Bremen | Temperate | EOS | | -0.22 | n.s. | -0.04 | |
| Brussels | Temperate | EOS | -0.07 | 0.13 | -0.02 | 0.00 | n.s. |
| Bucharest | Temperate | EOS | | 0.16 | | 0.04 | n.s. |
| Budapest | Temperate | EOS | | 0.07 | | 0.04 | n.s. |
| Bursa | Mediterranean | EOS | n.s. | n.s. | | -0.28 | |
| Copenhagen | Temperate | EOS | | 0.09 | n.s. | 0.02 | n.s. |
| Dresden | Temperate | EOS | n.s. | 0.30 | n.s. | 0.09 | |
| Dublin | Temperate | EOS | n.s. | n.s. | n.s. | 0.06 | |
| Edinburgh | Temperate | EOS | | n.s. | n.s. | 0.15 | |
| Eskişehir | Mediterranean | EOS | | 0.34 | | -0.26 | |
| Frankfurt | Temperate | EOS | n.s. | 0.12 | n.s. | n.s. | n.s. |
| Gaziantep | Mediterranean | EOS | | -0.14 | | | |
| Genoa | Temperate | EOS | n.s. | | n.s. | 0.11 | 0.04 |
| Glasgow | Temperate | EOS | n.s. | 0.08 | n.s. | 0.02 | |

| | | | | | | | |
|---|---|---|---|---|---|---|---|
| Göteborg | Temperate | EOS | 0.05 | -0.04 | 0.02 | 0.01 | |
| Hamburg | Temperate | EOS | n.s. | n.s. | n.s. | 0.01 | |
| Hannover | Temperate | EOS | n.s. | 0.11 | n.s. | n.s. | |
| Helsinki | Temperate | EOS | n.s. | 0.13 | n.s. | 0.05 | |
| Istanbul (MA) | Mediterranean | EOS | 0.16 | 0.14 | n.s. | -0.12 | n.s. |
| İzmir | Mediterranean | EOS | n.s. | -0.22 | | -0.32 | |
| Konya | Mediterranean | EOS | | 0.23 | | | |
| Kraków | Temperate | EOS | | 0.10 | n.s. | 0.07 | -0.06 |
| Leipzig (MA) | Temperate | EOS | | n.s. | n.s. | -0.22 | n.s. |
| Lisbon | Mediterranean | EOS | -0.83 | -0.16 | | -0.36 | |
| Liverpool | Temperate | EOS | | 0.18 | n.s. | 0.10 | |
| Łódź | Temperate | EOS | n.s. | 0.17 | 0.04 | 0.05 | n.s. |
| London | Temperate | EOS | n.s. | n.s. | n.s. | 0.19 | n.s. |
| Lyon | Temperate | EOS | | -0.20 | -0.37 | -0.08 | -0.13 |
| Madrid | Mediterranean | EOS | 0.33 | -0.02 | | 0.07 | |
| Málaga | Mediterranean | EOS | | 0.42 | | 0.31 | |
| Marseille | Mediterranean | EOS | n.s. | n.s. | 0.19 | n.s. | |
| Mersin | Mediterranean | EOS | | n.s. | | -0.37 | |
| Milan | Temperate | EOS | 0.56 | 0.03 | 0.11 | 0.01 | -0.02 |
| Munich | Temperate | EOS | | 0.18 | n.s. | 0.03 | |
| Naples | Mediterranean | EOS | -0.31 | 0.53 | n.s. | -0.04 | -0.09 |
| Nürnberg | Temperate | EOS | 0.10 | 0.43 | 0.04 | 0.13 | |
| Oslo | Temperate | EOS | 0.06 | 0.14 | n.s. | 0.08 | |
| Palermo | Mediterranean | EOS | 2.14 | n.s. | | n.s. | |
| Paris | Temperate | EOS | | 0.14 | 0.01 | 0.02 | 0.01 |
| Poznań | Temperate | EOS | n.s. | 0.17 | n.s. | n.s. | |
| Prague | Temperate | EOS | n.s. | 0.17 | n.s. | 0.05 | n.s. |
| Randstad (MA) | Temperate | EOS | | 0.27 | n.s. | 0.04 | |
| Riga | Temperate | EOS | n.s. | 0.03 | 0.09 | 0.03 | |
| Rin-Ruhr (MA) | Temperate | EOS | n.s. | 0.20 | -0.01 | 0.03 | n.s. |
| Rome | Mediterranean | EOS | n.s. | n.s. | n.s. | -0.14 | n.s. |
| Sarajevo | Temperate | EOS | | -0.18 | | 0.04 | -0.15 |
| Sevilla | Mediterranean | EOS | | -0.40 | | -0.43 | |
| Sheffield | Temperate | EOS | | 0.30 | -0.12 | 0.12 | n.s. |
| Skopje | Temperate | EOS | | 0.11 | | n.s. | |
| Sofia | Temperate | EOS | | 0.23 | n.s. | 0.08 | -0.13 |
| Stockholm | Temperate | EOS | 0.03 | 0.15 | 0.04 | 0.05 | |
| Stuttgart | Temperate | EOS | | 0.06 | n.s. | n.s. | n.s. |
| Toulouse | Temperate | EOS | | -0.43 | | -0.30 | |
| Turin | Temperate | EOS | | 0.08 | n.s. | 0.05 | n.s. |
| Valencia | Mediterranean | EOS | | -0.93 | | -0.42 | |
| Vienna | Temperate | EOS | 0.17 | -0.10 | 0.05 | 0.02 | -0.02 |
| Vilnius | Temperate | EOS | n.s. | 0.07 | n.s. | 0.01 | |
| Warsaw | Temperate | EOS | n.s. | 0.04 | 0.01 | 0.02 | 0.16 |
| Wrocław | Temperate | EOS | | 0.11 | n.s. | 0.04 | 0.16 |
| Zagreb | Temperate | EOS | | n.s. | | 0.07 | -0.13 |
| Zaragoza | Mediterranean | EOS | | -0.27 | | n.s. | |

**Appendix D.** Phenological response to year (a) and elevation (b) of the main vegetation types expressed as the regression coefficients ($\beta_Y^P$ and $\beta_E^P$) obtained from multiple regression models using urbanisation, year and elevation as predictors.

The average SOS of the vegetation types advanced 4.68 ± 2.83 days (mean ± sd) and the LOS was extended 3.8 ± 2.33 days in the last two decades. The LOS of deciduous, evergreen and mixed forests and sparse woody vegetation has increased by 0.15-0.30 days/year over the last two decades. We also found an average positive relationship between elevation and SOS for all the vegetation types considered (0.30-34.17 days/km). The relationship between elevation and the other two phenological indexes (LOS and EOS) was highly dependent on the type of vegetation considered.

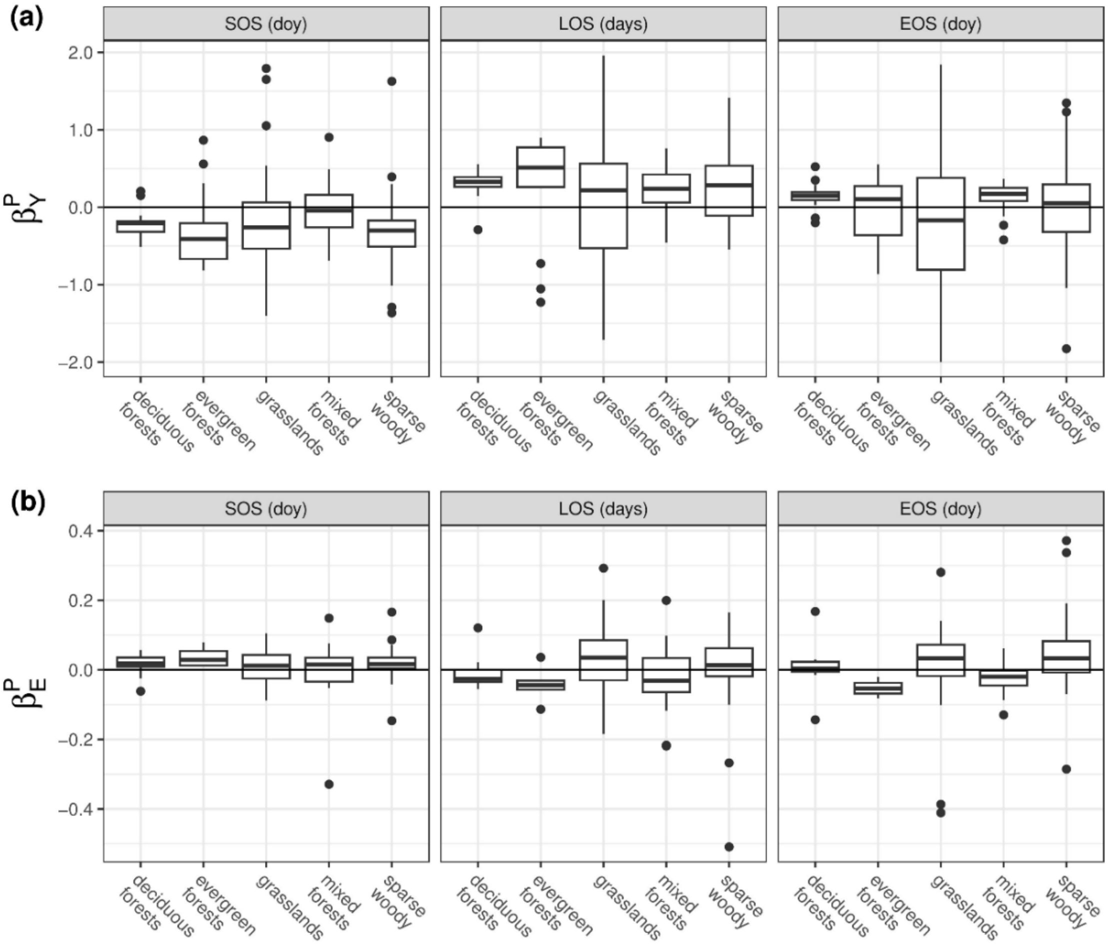

**Appendix E.** Partial effect of urbanisation intensity on the phenological indexes grouped by macrobioclimate. For each metropolitan area, random forest models were performed to predict the general phenological response to urbanisation. The phenological indexes were normalised by subtracting the mean and dividing by the standard deviation. SOS: start of season date, LOS: length of season, EOS: end of season date.

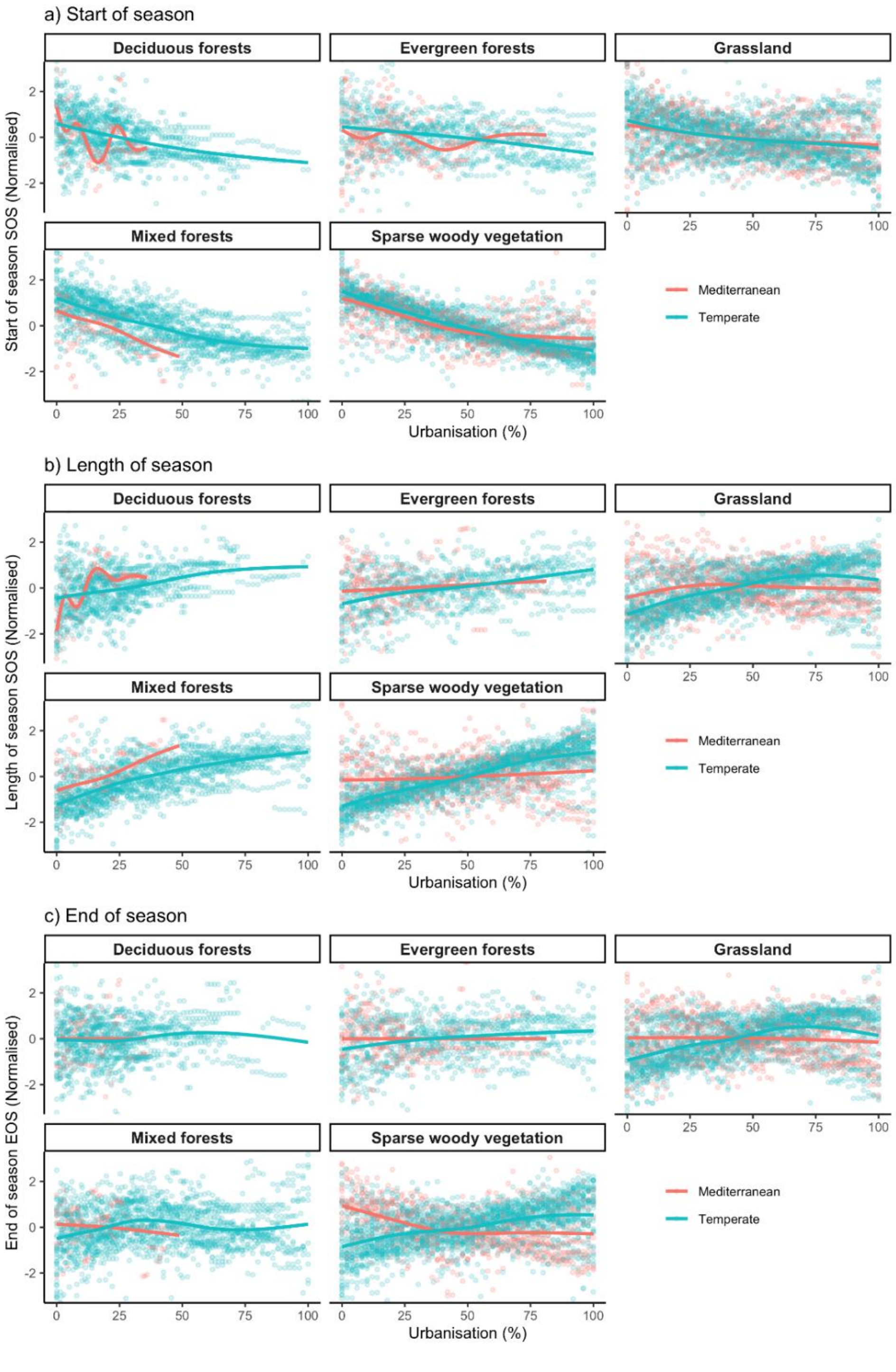

**Appendix F.** Relationship between land surface temperature (LST) and urbanisation intensity ($\beta_U^{LST}$) for each metropolitan area, expressed as a function of annual mean temperature. We used the average day and LST per season as the response variable, and ran multiple linear regressions using urbanisation intensity, year and elevation as explanatory variables. DJF: winter, MAM: spring, JJA: summer, SON: autumn.

LST increased with urbanisation across European metropolitan areas (Figure 6a). The rate of increase in LST with urbanisation ($\beta_U^{LST}$) decreased with annual mean temperature and was greater in Temperate than in Mediterranean metropolitan areas during both day and night. $\beta_U^{LST}$ in Temperate metropolitan areas was greater during spring (MAM) and summer (JJA). LST has increased over the last two decades across European metropolitan areas (Figure 6b). Metropolitan areas in the Temperate macrobioclimate showed an especially sharp rise in winter temperatures (DJF).

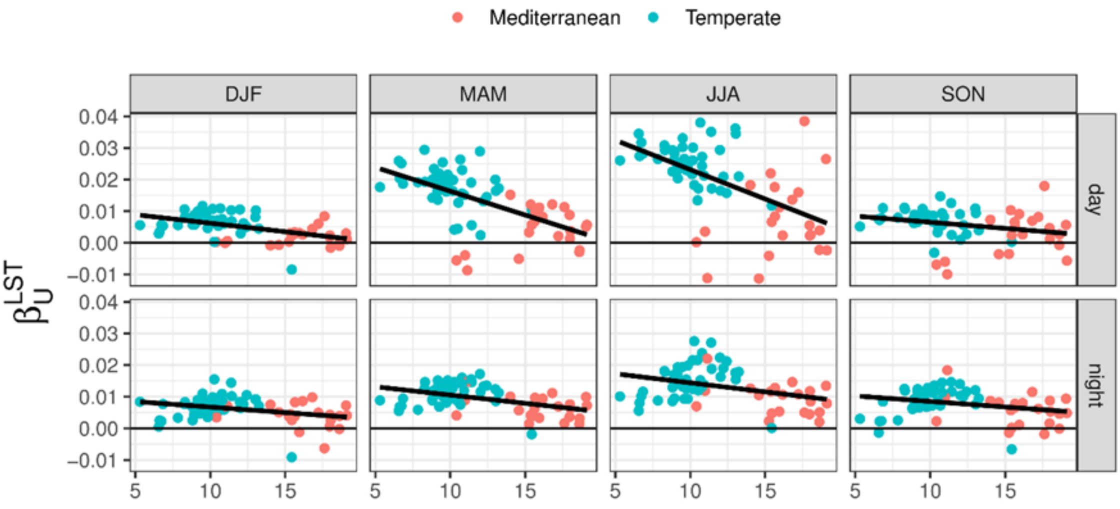
